# The Parasite Intraerythrocytic Cycle and Human Circadian Cycle are Coupled During Malaria Infection

**DOI:** 10.1101/2022.08.02.499203

**Authors:** Francis C. Motta, Kevin McGoff, Robert C. Moseley, Chun-Yi Cho, Christina M. Kelliher, Lauren M. Smith, Michael S. Ortiz, Adam R. Leman, Sophia A. Campione, Nicolas Devos, Suwanna Chaorattanakawee, Nichaphat Uthaimongkol, Worachet Kuntawunginn, Chadin Thongpiam, Chatchadaporn Thamnurak, Montri Arsanok, Mariusz Wojnarski, Pattaraporn Vanchayangkul, Nonlawat Boonyalai, Philip L. Smith, Michele Spring, Krisada Jongsakul, Ilin Chuang, John Harer, Steven B. Haase

## Abstract

During infections with malaria parasites *P. vivax*, patients exhibit rhythmic fevers every 48 hours. These fever cycles correspond with the time parasites take to traverse the Intraerythrocytic Cycle (IEC) and may be guided by a parasite-intrinsic clock. Different species of *Plasmodia* have cycle times that are multiples of 24 hours, suggesting they may be coordinated with the host circadian clock. We utilized an *ex vivo* culture of whole blood from patients infected with *P. vivax* to examine the dynamics of the host circadian transcriptome and the parasite IEC transcriptome. Transcriptome dynamics revealed that the phases of the host circadian cycle and the parasite IEC were correlated across multiple patients, suggesting that the cycles are coupled. In mouse model systems, host-parasite cycle coupling appears to provide a selective advantage for the parasite. Thus, understanding how host and parasite cycles are coupled in humans could enable anti-malarial therapies that disrupt this coupling.

## Introduction

Malaria disease in humans, caused by infection with the parasite genus *Plasmodium*, remains a global health crisis with an estimated 241 million cases and 627,000 deaths worldwide in 2020 (World Health, 2021). Effective treatment is complicated by continued development of drug resistance, already established in *P. falciparum* and emerging in *P. vivax* (Kozlov, 2021; Price et al., 2014; Wellems and Plowe, 2001). Drug resistance, combined with the threat of zoonotic jumps to humans by simian-infecting parasites (Bykersma, 2021), is motivating researchers to better understand fundamental *Plasmodium* biology in the hopes of revealing novel drug targets and innovative and improved methods of treatment (Cowell and Winzeler, 2019; Ippolito et al., 2021).

The multistage lifecycle of *Plasmodium* includes a distinct intraerythrocytic developmental cycle (IEC) inside the host organism that is characterized by periodic rupture, egress and (re)invasion of host erythrocytes (red blood cells [RBCs]). While inside an RBC, *Plasmodium* is observed to progress through morphologically distinct stages ending with schizont stage, during which the parasite undergoes multiple rounds of asexual division. The resulting merozoites then rupture the RBC and rapidly infect new RBCs, beginning the cycle anew.

For human-infecting *Plasmodium* species, the infecting parasite populations synchronously advance through the IEC, maintaining precise periods of 24, 48, or 72 hours depending on the species. Recent evidence suggests that the maintenance of synchronized progression through the *Plasmodium* IEC, and thus the synchronized bursting of RBCs and subsequent release of parasites is controlled by parasite intrinsic molecular oscillators that may interact with host circadian cycles (Rijo-Ferreira et al., 2020; Smith et al., 2020).

Synchronized bursting on its own may provide a benefit to the infecting parasite populations - either by overwhelming the host immune system (McQueen and McKenzie, 2008) or through evasion of the immune response (Rouzine and McKenzie, 2003). Furthermore, alignment of the synchronized IEC to the host circadian rhythm could further benefit the parasite by avoiding peaks of some immune activity that may be under circadian control (Ikuta and Scheiermann, 2020; Mideo et al., 2013), such as rhythmic fluctuation of immune cells in blood (Yuan et al., 2020). Alternatively, parasites may time their reproduction to maximize access to host resources (Hirako et al., 2018; Prior et al., 2018). This alignment could help explain the apparent reduced parasite fitness induced by misalignment of IEC stage and host circadian phase (Mideo et al., 2013; O’Donnell et al., 2011; Reece et al., 2017). However, whether the Plasmodium IEC aligns with the host circadian rhythms in humans is unknown.

Does the *Plasmodium* IEC align with the host circadian rhythms in humans? In this study, we examine the alignment of transcriptional phases of the *P. vivax* IEC to human host circadian rhythm across a cohort of 10 human-parasite pairs. Using high-resolution time-series transcriptomics to measure both host and parasite dynamics, we observe significant correlation between the phases of the parasite IEC and the host circadian rhythm. These findings provide evidence that the timing of the IEC in human-infecting *Plasmodium* parasites is aligned with a host circadian signal detectable in whole blood *ex vivo*.

## Results

### *G*ene expression dynamics of host and parasite cycles in cultured whole blood from infected hosts exhibit rhythmic behavior

19 adult participants (anonymized as participants 01–19) voluntarily presented at clinics in Na Chaluai and Buntharik Districts in Ubon Ratchathani Province, Thailand seeking treatment for suspected malaria infection. Prior to treatment, each participant underwent screening for study enrollment (see **STAR Methods** for study design and inclusion criteria details). Following the initial screening, 10 of the 19 participants (participants 02, 08, 09, 10, 11, 13, 16, 17, 18, 19) were enrolled in Part B and whole blood samples were collected and further divided into 16 samples (Figure 1A). Other participants exhibited insufficient parasitemia levels or did not meet all inclusion/exclusion criteria. All eligible participants’ parasite populations were determined by microscopy to be well synchronized (greater than 80%) in the early trophozoite developmental stage at the time of the screening (Table S3 and Figure 1B).

**Figure 1.**
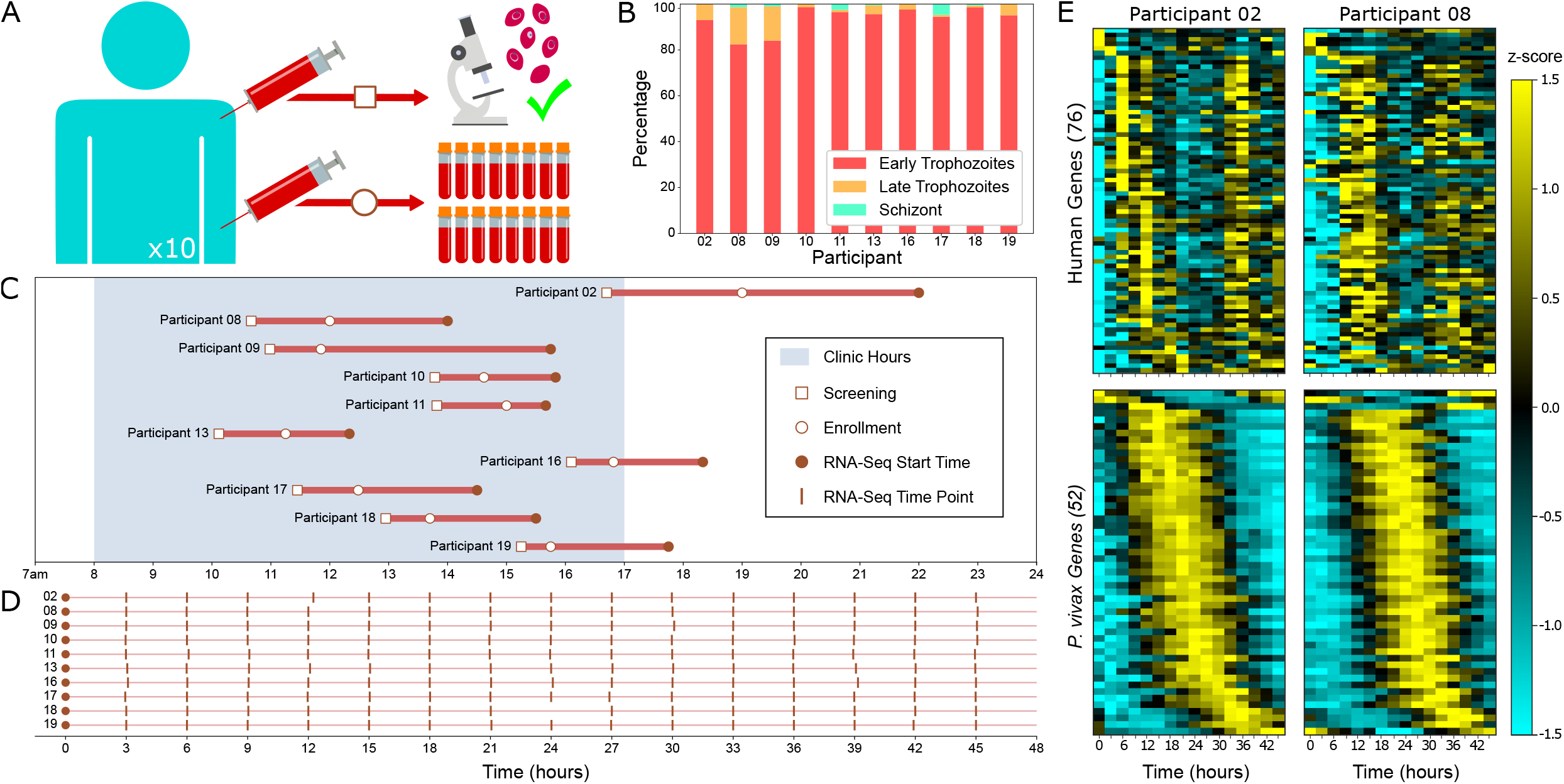
Experimental protocol and data collection: (A) Diagram of the clinical study sampling design. Each participant underwent a single blood draw that is used for screening to ensure uncomplicated mono-infection by *P. vivax*, and a second draw used for RNA sequencing. If the microscopy screening showed asexual parasitemia ≥0.1% and hematocrit ≥25%, and parasites were observed to be well-synchronized in early or late trophozoite phase, the participant was enrolled and a second blood draw was taken and divided into 16 parts for subsequent RNA-Seq analysis. (B) Times of day at which each study participant was screened and enrolled, and the time of day of the first time point in the RNA-Seq time series experiment. (C) For each participant, the times relative to the start time at which each of the 16 samples were aliquoted and frozen for later RNA sequencing. (D) *P. vivax* developmental stage percentages at the screening time for each participant. (E) Representative z-score heatmaps of genes which exhibit periodic expression patterns at a specified period (24 hours for human genes and 48 hours for *P. vivax* genes) in participants 02 and 08. For each participant and each organism, periodic genes are taken to be those whose maximum expression across the time series is at least 1 TPM and which exhibit at least the same degree of periodicity as the top 5% of genes when ranked by the JTK-Cycle periodicity score. Genes are consistently ordered in each pair by participant 02’s peak expression time during the first period. See also Figure S1 and Figure S2.

Participants entered the clinics, were screened, and enrolled across a wide range of times of day (Figure 1C). After a delay following collection (Figure 1C), each of the 16 whole blood aliquots were initially processed (**STAR Methods**) and then frozen roughly every 3 hours over 45 hours (Figure 1D). All but one sample (time point number 9 corresponding to hour-24 for participant 18) survived for subsequent RNA sequencing (Figure 1D). Reads from RNA sequencing of the remaining 159 samples were aligned to both human and *P. vivax* transcriptomes, quantified using TPM normalization and minimally post-processed (**STAR Methods**) to produce 10 pairs of 16-time-point time series datasets capturing *ex vivo*, whole transcriptome abundance profiles for human participants and their *P. vivax* populations over 45 hours. Read mapping percentages reported by the RNA sequencing read alignment are provided in Figure S1.

Focusing on rhythmic transcripts that are common to participant pairs, we observed a high degree of similarity in the ordering of peak expression and the dynamics of expression across all pairs of parasite IEC transcriptomes and many pairs of human circadian transcriptomes (Figures 1E and S2). Also, the periodic transcript programs of parasites and humans appear qualitatively to be largely well-aligned. For example, subsets of rhythmic genes common to both participant 02 and 08 appear by eye to be consistently ordered in both the human and the *P. vivax* populations (Figure 1E). Moreover, there is a noticeable shift forward in expression peaks of participant 08 compared with 02, in both the parasite sample and the human sample. This suggests participant 02’s host and parasite transcriptional phases were both slightly ahead of participant 08’s. These qualitative phase shifts are apparent across many pairs in which heatmaps show enough consistency in gene order to make a qualitative assessment of the relative transcriptional phases by visual inspection (Figure S2). Analogous figures—with each of the other participants serving as the reference—can be generated using the data and code repositories associated with this manuscript.

### Gene expression dynamics in *ex vivo* whole blood samples are similar to previous *in vivo* studies

The *ex vivo* time series is a novel approach to examining both host and parasite rhythms by transcriptomic analysis. To verify that this approach was producing results similar to those observed previously in studies examining only host *or* parasite rhythms, we quantitatively compared our data to earlier studies.

Periodic transcription of temporally ordered genes in the IEC has been confirmed in previous studies across different malaria species (Bozdech et al., 2008; Smith et al., 2020). Other studies involving *P. vivax* used an *ex vivo* system in which white blood cells were removed by filtration and utilized a microarray platform for measuring transcriptome dynamics rather than an RNASeq platform utilized in this study (Bozdech et al., 2008). Thus, these other studies were unable to simultaneously assess both parasite and host rhythms.

First, we compared *P. vivax* transcript dynamics in this study to the dynamics of three *P. vivax* isolates published in (Bozdech et al., 2008). While phase-shifted 3-6 hours from the (Bozdech et al., 2008) *P. vivax* isolates, all ten participants displayed highly similar transcript dynamics to all three isolates (Figure 2A). Temporal ordering of highly periodic genes appears to be conserved between the *P. vivax* in our participants and the (Bozdech et al., 2008) *P. vivax* isolates (Figure 2B). Interestingly, Participant 09, appears phased almost halfway through the IEC cycle ahead of the other nine participants (Figure 2A), even though Participant 09’s developmental stage agrees with the other nine participants (Figure 1D). Temporal ordering of highly periodic genes appears to be conserved between our ten participants and the (Bozdech et al., 2008) *P. vivax* isolates (Figure 2B). In both studies, peaks of gene expression are evenly distributed throughout the IEC and the beginning of a second wave of gene expression, indicated by the increasing gene expression of early phased IEC genes later in the time series, can be seen clearly (Figure 2B). The gene expression of some late IEC phased genes seems to appear at the beginning of the time series in our participants (presumably from the previous cycle), which is perhaps less visible in the (Bozdech et al., 2008) isolates (Figure 2B).

**Figure 2.**
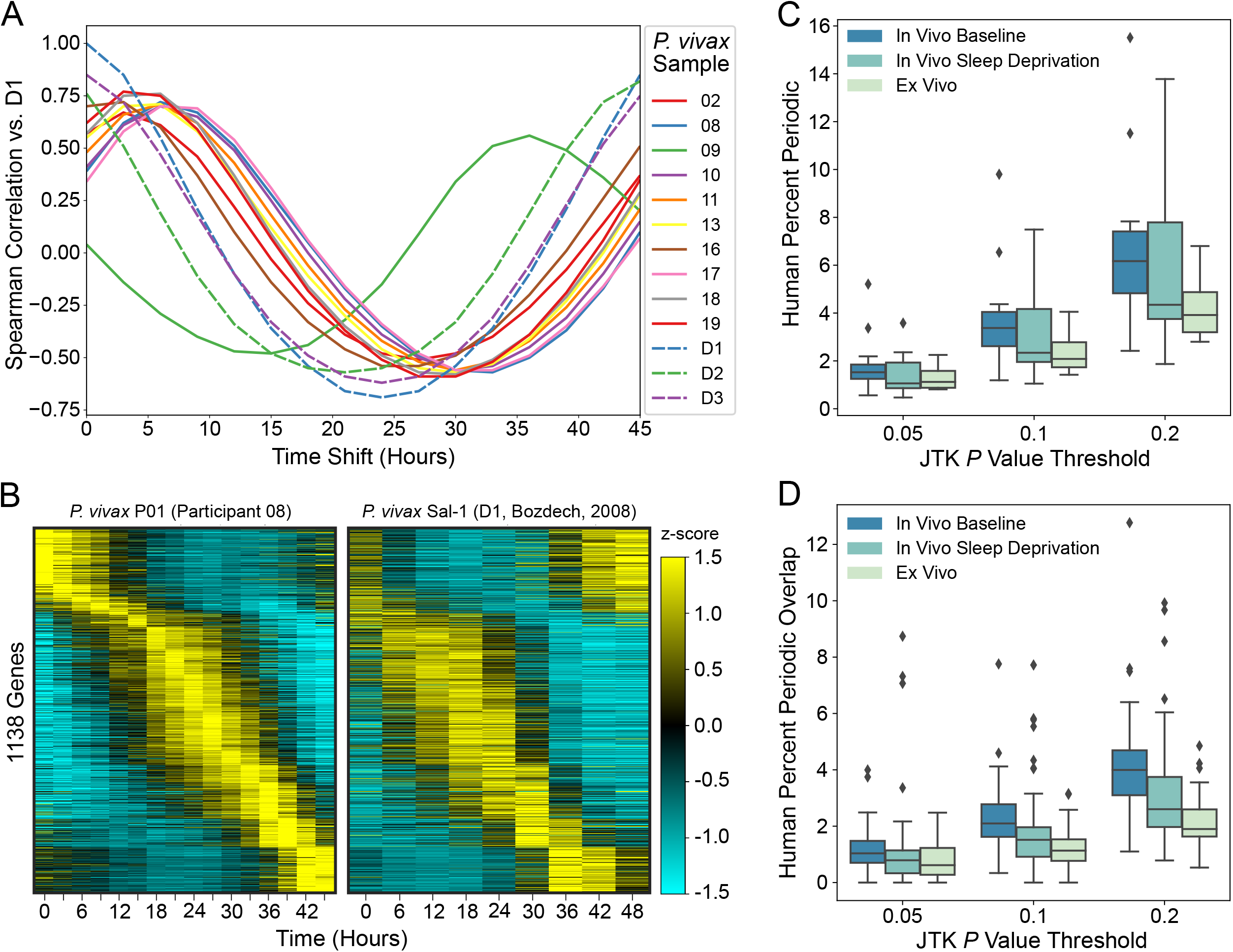
Comparisons of RNA-Seq time series data to earlier studies: (A) The average pairwise Spearman correlation between one of the *P. vivax* (Sal-1) isolates from the *ex vivo* time series RNA-Seq experiment reported in (Bozdech et al., 2008), the other two samples reported in that study, and each of the 10 participants in this study. Correlations are the average over all time points shifted by the specified amount, with wrapping assuming a 48-hour period. (B) z-score heatmaps of highly periodic (JTK *p* value ≤0.005) *P. vivax* (P01) genes in participant 08 that also appear in sample D1 from (Bozdech et al., 2008). All heatmaps contain 1138 genes and are ordered by the time of peak expression in participant 08. Analogous plots can be generated for the other participants using data and code repositories associated with this manuscript (Key resources table). (C) Box-and-whisker plots comparing the average percentage of expressed genes common to the three conditions and achieving the specified JTK *p* value thresholds. (D) Box-and-whisker plots comparing the average percent overlap among the rhythmic genes achieving the specified JTK *p* value thresholds. Distributions in (C) represent either the 14 participants in the *in vivo* study, (Arnardottir et al., 2014), under baseline or sleep deprivation conditions, or the 10 participants in this study. Distributions in (D) represent all participant pairs within each study. In (C) and (D) boxes indicate distribution quartiles and whiskers extend from minimum sample values to samples at most 1.5 times the interquartile ranges, with values outside this range plotted as outliers.

Previous studies examining circadian gene expression in human whole blood have found the circadian signal to be weaker than that found in other tissues in which many genes are known to be highly rhythmic (Arnardottir et al., 2014; Möller-Levet et al., 2013). We also expect factors related to *ex vivo* sampling in our study could contribute to degradation of circadian signal. For example, our samples were collected *ex vivo*, and so all environmental entrainment signals (e.g., light/dark cycles and eating) have been removed, which may contribute to a reduced circadian signal as natural variability in oscillation periods will cause synchrony loss across a population or cells (Leise et al., 2012). Additionally, our study covers two circadian cycles (two 24-hour periods), which can further dampen the circadian signal, due in part to synchrony loss. Finally, it is not clear that human peripheral blood mononuclear cells would remain healthy in *ex vivo* culture, which could certainly affect the circadian rhythm or the integrity and stability of transcripts.

In addition to *ex vivo* conditions, a variety of lifestyle and environmental variables encountered by participants in this study could impact the rhythmic transcriptomic signals observed for human subjects in the study. Some of the participants are night shift workers while others reported sleep disruptions (Table S3), both known factors that can impact circadian rhythmicity in blood (Arnardottir et al., 2014; Möller-Levet et al., 2013). Also, we cannot rule out the impacts, direct or indirect, that malaria infection may have on a person’s circadian rhythm.

To estimate the cumulative effect of the above issues on the detection of circadian rhythmicity, we compared our participants’ circadian transcriptomes to a controlled study that investigated the impacts of sleep deprivation on the circadian transcriptome in human whole blood (Arnardottir et al., 2014). We found only a moderate decrease in the number of circadian transcripts found in our participants compared to participants under normal sleep conditions (Figure 2C). This moderate decrease in circadian signal in our participants compared to *in vivo* data under normal sleep conditions is comparable to those seen in participants with sleep deprived conditions compared to normal sleep conditions. In particular, the median percentage of rhythmic genes in the *in vivo* sleep study under sleep deprivation conditions dropped by between 0.46% and 1.82% across the range of thresholds, while our *ex vivo* study observed the percentage of rhythmic genes drop by between 0.4% to 2.24% across the same thresholds (Figure 2C). Additionally, the confounding factors in our study appeared to have had a similar effect on the expression of the circadian transcriptome across the ten participants that sleep deprivation had on the participants in the sleep deprivation study. The median percentage of common rhythmic genes out of the combined rhythmic genes among pairs of participants in the *in vivo* sleep study under sleep deprivation conditions dropped between 0.25% to 1.39% across the range of thresholds, while our *ex vivo* study saw this percentage of rhythmic gene overlap drop from 0.42% to 2.1% across the same thresholds (Figure 2D). Our results suggest that the circadian transcriptional program observed in the whole blood of ten participants in this study is largely intact, but it may be partially degraded by a combination of internal and external factors.

### Relative phasing of *P. vivax* IEC cycles are largely similar across hosts

Based on microscopic analyses of blood samples before the start of the transcriptome time series, we expected the parasites to be in a similar phase of their IEC in each of the 10 subjects. However, the lag between microscopic analysis and the start of the transcriptomic analyses varied between subjects (Figure 1C). To compare the relative phasing of the transcriptome analyses, we focused the analysis on genes expressed at the end of the IEC before egress of the parasites (see Methods) (Figure 3A). All but one of the participants exhibit late IEC gene expression programs just prior to the end of the time series (between 36 and 45 hours), suggesting the IECs across subjects are in approximately the same phase.

**Figure 3.**
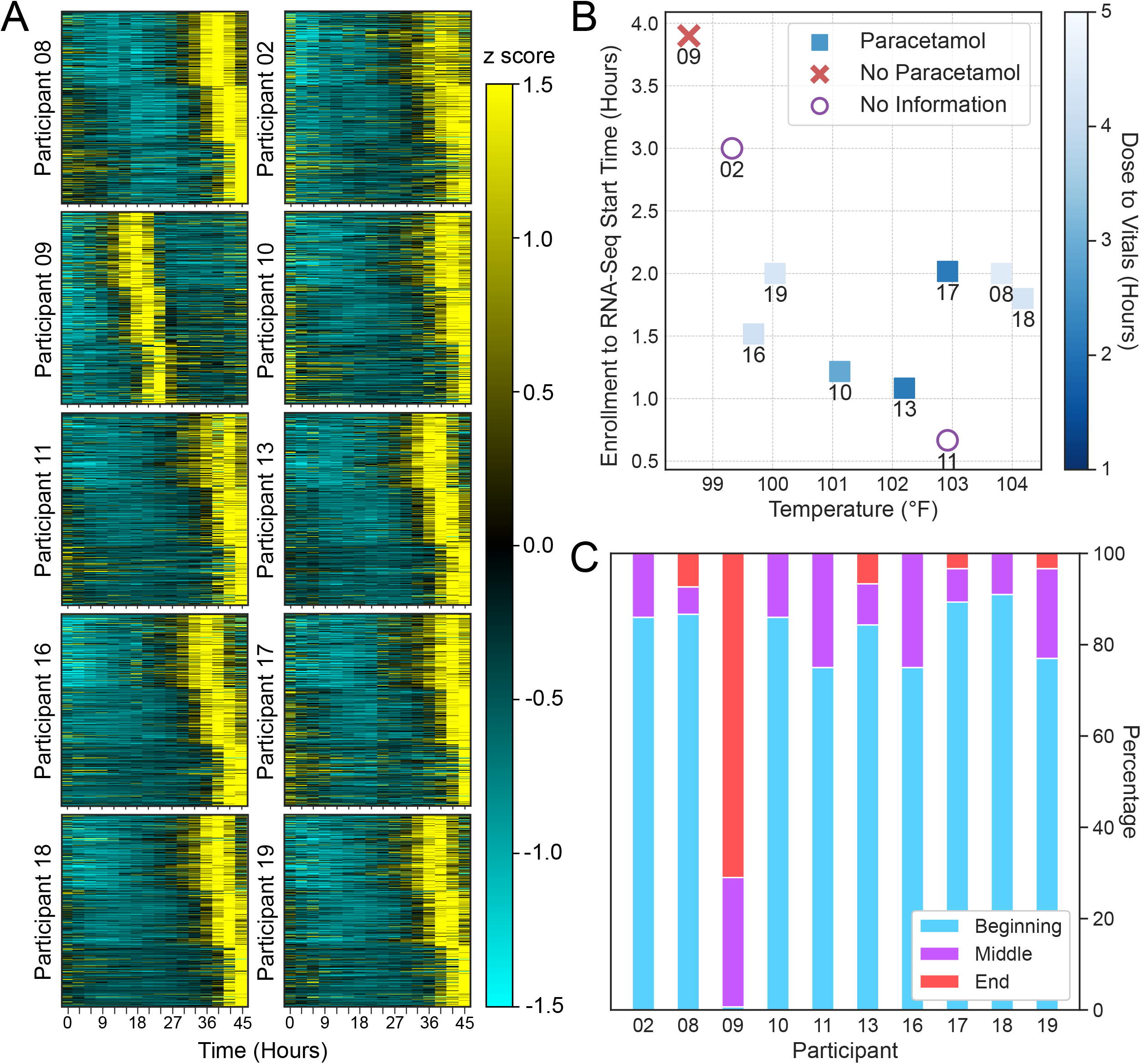
Evidence of participant 09 P. vivax transcriptional phase shift: (A) Z-score heatmaps of highly periodic genes (JTK *p* value ≤0.005) *P. vivax* genes which show peak expression in one of the final three timepoints (i.e. times greater than 36 hours) for participant 08. Each participant’s heatmap shows the subset of genes with non-zero expression, ordered according to the peak expression in participant 08. (B) Human participant metadata: body temperature plotted against the duration between the time of the enrollment blood draw and the RNA-Seq time series start time, decorated by if and when a fever reducer was administered relative to when the vitals were taken, if known. (C) Refined *P. vivax* stage percentages grouped into beginning, middle, and end of the early trophozoite stage. See also Figure S3 and Table S3.

Participant 09 is an outlier, and the IEC appears to be phase-shifted from the others by 18-21 hours. In support of this shift, the timepoint-wise Spearman correlations between *P. vivax* transcriptome expression profiles in this study and the data reported previously (Bozdech et al., 2008) show peak correlations between isolate D1 and participant 09 occurring at a relative shift of around −12 hours, while peak correlations with the other 9 participants in this study occur between relative hour 3 and 6 (Figure 2A). Moreover, analogous plots of early peaking and middle peaking genes in a chosen reference participant also show consistent shifts in participant 09 (not shown) and demonstrate that despite the substantial phase-shift, dynamic gene expression could be observed throughout the entire IEC in participant 09.

These findings suggest that the parasites from participant 09 were phased in the latter stages of trophozoite phase when the time series began rather than the early stages. Findings also suggest that at least part of the transcriptional program continues following what ostensibly corresponds to *ex vivo* egress from RBCs of 09’s parasites. This is expected if there exists an autonomous transcriptional regulatory network controlling the timing and ordering of expression during the *P. vivax* IEC as has been shown in recent studies (Rijo-Ferreira et al., 2020; Smith et al., 2020). It is not known whether parasites re-infected RBCs in the whole blood after egress halfway through the time series so it is possible that the transcriptional rhythm continues in parasites that have not reinvaded.

Additional evidence is consistent with the observation that participant 09’s *P. vivax* transcriptional phase is advanced by approximately 40% of its developmental cycle relative to other participants. First, participant 09 had the lowest body temperature of any participant, and in fact did not enter the clinic with a fever as others had. Notably, Participant 09 had not taken a fever reducer prior to vitals being taken, while the other participants are known to have taken a fever reducer within 5 hours of the time at which vitals were taken, except for participants 02 and 11 for which that information is not known (Figure 3B and Table S3). These observations suggest 09’s parasites were phase advanced relative to the other participants at the start of their time series, given that egress and reinvasion corresponds to the sharply-delineated, short-duration (<8 hour) fever spikes in *P. vivax* infections (Karunaweera et al., 1992).

Participant 09 also had the longest duration among all participants between their study enrollment blood draw and the processing of their hour-0 sample, providing the parasites a longer duration than other samples to progress through the IEC before the start of the RNA sequencing experiment (Figure 1C, Figure 3B, and Table S3). Finally, microscopists performed a follow-up evaluation of Study B participant blood slides to quantify developmental stage according to a refined categorization of the *P. vivax* early trophozoites into beginning, middle, and end phases based on morphological characteristics as illustrated in Figure S3. This reanalysis found that 09’s parasites had mostly progressed to late-phase early trophozoites, while all other participant parasites were primarily in early-phase early trophozoites (Figure 3C), which is consistent with 09’s advanced transcriptional phase relative to the other participants.

### *P. vivax* IEC phase and host circadian phase are correlated

If there is an alignment between the *P. vivax* IEC phase and the host circadian phase, due to the 2-to-1 coupling between the 48-hour IEC and 24-hour circadian rhythms, then there necessarily will be two IEC phases corresponding to this host phase, (time point 0 and time point 24 in Figure 4A). Thus, it is not surprising that among hosts with similar circadian phases we observe a parasite population which has progressed roughly halfway through its IEC cycle relative to the other parasites. In this way, participant 09 is not an outlier, but rather confirms expectation.

**Figure 4.**
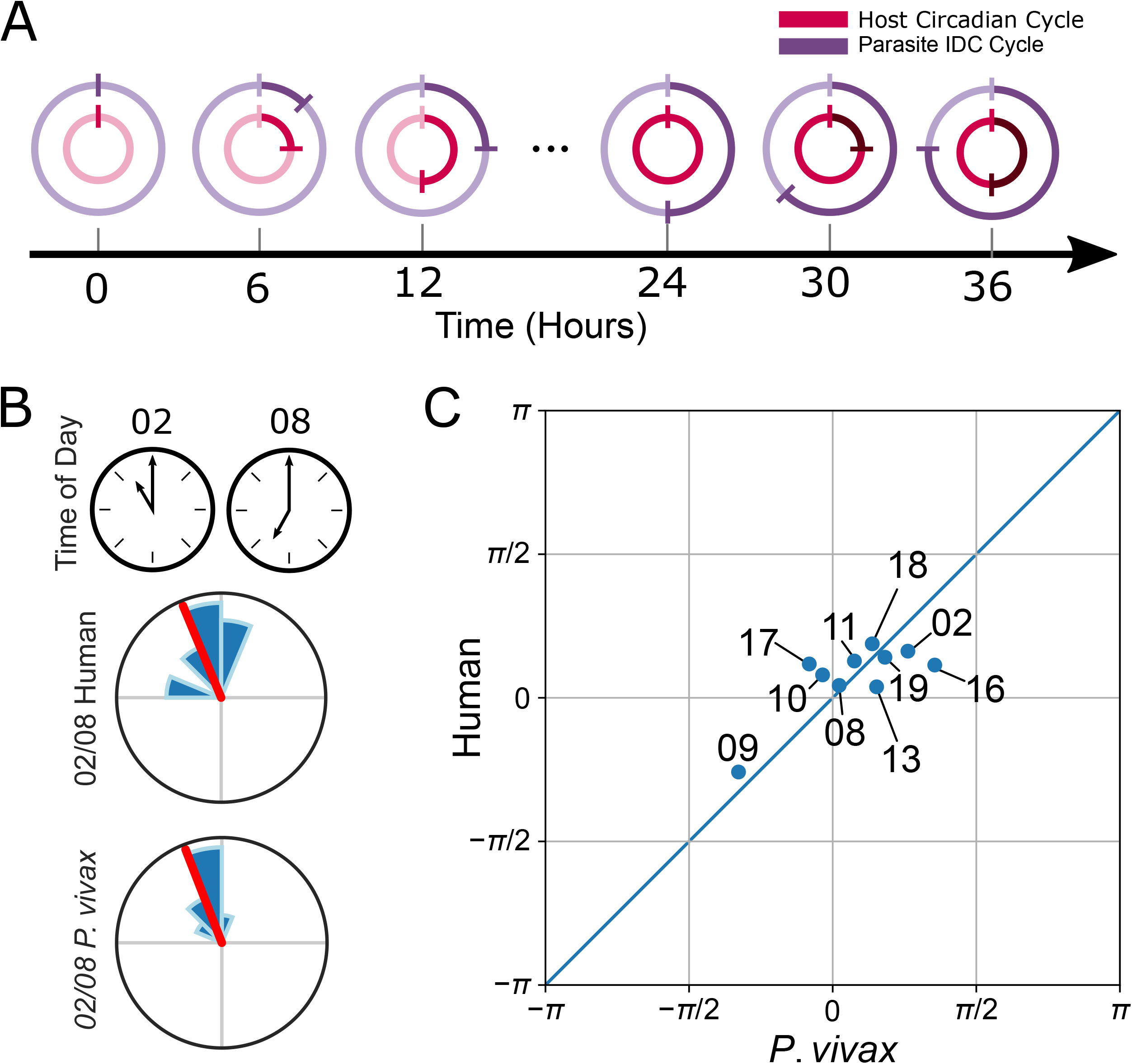
Human phase, *P. vivax* phase, and wall time circular correlations: (A) Cartoon illustrating the correspondences between elapsed time and the phases of a 48-hour cycle and a 24-hour cycle, indicating the difference between the circadian cycle phase and twice the intraerythrocytic developmental cycle phase is expected to be constant over time. (B) The times of day on 24-hour clocks of the first time points in the RNA-Seq time series (top) and the distributions of gene phase differences estimated from the overlapping periodic human (middle) and *P. vivax* (bottom) genes for participants 02 and 08. The radii of the histogram slices have been scaled so the area of each slice is proportional to the number of data points it represents. (C) Scatter plots of inferred human and *P. vivax* phases. For *P. vivax*, all phase difference estimates were first multiplied by two and reduced modulo 2*π* before visualization. All phases are shifted to the interval [–*π,π*) radians and the 0 phases were chosen to be the circular mean human phase. The left and right edges and the top and bottom edges should be identified since the phase/angle -*π* is equal to *π* on a circle. See also Figure S4.

Recent studies demonstrate that an intrinsic clock controls the IEC of some malaria parasite species (Rijo-Ferreira et al., 2020; Smith et al., 2020). In studies involving mice-infecting *Plasmodium* species with 24-hour IECs, their cycles appeared to be well aligned with the 24 hour circadian cycle of the host (O’Donnell et al., 2011; Rijo-Ferreira et al., 2020), and that alignment of parasite and host cycles was observed to be important to maintain parasite fitness. Here we aimed to determine whether a similar alignment of the *P. vivax* 48-hour IEC could be observed with the host circadian clock in humans.

By visual inspection, the apparent timings of peak expression in rhythmic *P. vivax* genes show notable qualitative shifts between pairs of participants, with the largest shift between 09 and the others (Figure 3A). We asked whether a phase shift in *P. vivax* IEC rhythms correlated with a shift in the host circadian rhythm. To address this question, we developed a data-driven pipeline to estimate the relative circadian transcriptional phase shifts between human participants and, separately, the relative IEC transcriptional phase shifts between their host parasites (**STAR Methods**).

If a *P. vivax* parasite, with its 48-hour IEC, has aligned a particular phase of its cycle (e.g., the egress from host RBCs) to a particular circadian phase in the host, with a 24-hour circadian cycle, then there would be a constant phase difference between the circadian phase of the host and twice the parasite IEC phase, as illustrated in Figure 4A. Since *P. vivax’s* expected IEC period length is twice as long as a host’s expected circadian cycle period, the time rate of change of the circadian phase should be twice the rate of change of the IEC phase. Therefore, if there is a parasite-host phase alignment, then the circadian phase difference between two hosts would be well correlated with twice the IEC phase difference between their parasite populations.

To estimate the transcriptional circadian phase difference between each pair of human participants, and separately the transcriptional IEC phase between their parasite populations, we computed the circular-mean phase difference between subsets of human and *P. vivax* genes that were highly oscillatory (at 24 and 48 hours, respectively) and highly expressed in both samples (Figure 4B and Figure S4A-B). This yielded 45 pairwise circadian phase difference estimates for human samples and 45 pairwise IEC phase difference estimates for *P. vivax* samples (Figure S4A-B).

Naturally, the pairwise phase shift estimates suffer from global inconsistencies. For example, the signed pairwise phase difference estimates from samples 02 to 11 and 11 to 19 imply a clockwise ordering of phases 02→11→19→02. On the other hand, taking the 11-to-19 and 02-to-19 phase shift estimates implies a clockwise ordering of 02→19→11→02. Such inconsistencies can be partially explained by the fact that each pairwise intersection of periodic genes may have different numbers and identities of genes, and so it is expected that some pairwise phase difference estimates will be less reliable than others (Figure S4C).

To correct errors in the pairwise phase estimates and compute a single circadian phase estimate for each human sample and a single IEC phase estimate for each *P. vivax* sample, we independently applied a phase synchronization method (Singer, 2011) to each collection of 45 pairwise phase estimates in hosts or parasites. Briefly, phase synchronization determines a globally consistent choice of phase variables that best explain a collection of noisy pairwise phase differences. Continuing with the above example, in the context of all pairwise phase estimates, the 02-11 estimate appears unreliable (Figure S4C). Specifically, after phase synchronization, the estimated phase difference between 02 and 11 changed from −0.5 to only −0.0171398, while the 02-to-19 and 11-to-19 phase differences were adjusted only marginally (0.023596 to −0.0106025 and 0.0528268 to 0.00653737 respectively). The original 02-to-11 pairwise phase difference estimate was based on only 3 genes whose apparent phase differences were highly disparate (Figure S4B), which may explain why the 02-11 pairwise phase estimate was discordant with other phase estimates. Unlike the pairwise circadian phase estimates, the *P. vivax* pairwise phase estimates were already remarkably globally concordant, exhibiting a high confidence circular correlation of 0.996461 (p value: 0.000004) with the pairwise phase difference estimates computed after phase synchronization (Figure S4C).

Importantly, our method of computing sample phase differences is entirely data-driven and depends on limited assumptions. For example, it makes no assumptions about which transcripts will reflect a circadian phase in whole blood during malaria infection, and instead it only assumes that the relative transcriptional phases in each organism will be most reliably estimated by subsets of highly expressed genes oscillating with the correct period. Moreover, this process uses multiple subsets of overlapping gene sets to derive globally consistent estimates of phases that best explain the pairwise phase differences seen in the data, some of which will undoubtedly be less reliable than others.

To compare the relative phase differences and identify the degree of alignment between the host circadian and parasite IEC phases, we computed the circular correlation ((Jammalamadaka and Sarma, 1988), **STAR Methods**) of the 10 pairs of estimated phases and found a circular correlation of 0.70 (p value: 0.019). The significance of the correlation was estimated using an exhaustive permutation test (Figure S4D, **STAR Methods**). The circular correlation between two random circular variables is 1.0 if and only if they differ by a constant, and it is invariant under a constant phase shift of either sample. The circadian and IEC phase pairs are plotted in Figure 4C, in which proximity to the diagonal indicates the degree of agreement between the relative phases of the hosts and their parasites. The significant shift in participant 09’s estimated IEC phase relative to the other parasite populations agrees with the by-eye estimate of 09’s relative phase advance based on the heatmaps shown in Figure 3A and the timepoint-wise correlation analysis shown in Figure 2A. To see this, observe that 18 hours in a 48 hour cycle is 6*π*/8 radians, which when multiplied by 2 yields a relative advance of 3*π*/2 radians, which is equivalent to a relative delay of -*π*/2 radians between 09 and the other participants, as seen in Figure 4C. Notably, twice the inferred 09 *P. vivax* phase is in excellent agreement with the inferred 09 human phase (Figure 4C).

Interestingly, the correlation between the estimated circadian phases and the time of day of the first time point in the RNA-Seq time series was found to be only 0.31 (p value: 0.178). To try and control for any underlying differences in the circadian phases of participants, we additionally computed correlations between circadian transcriptional phases and the phases determined by the times between each participant’s first RNA-Seq measurement and their self-reported usual wake-up times as well as their self-reported recent wake times (Table S3) and found the correlations did not improve (Table 2).

By restricting to the highest amplitude and most robustly oscillating genes, we worried that there may be too few genes in the pairwise intersections of human samples to provide reliable estimates of circadian phases. Thus, to assess the sensitivity of our conclusions to the choice of periodicity and activity thresholds, we considered a wide range of thresholds, dramatically relaxing them in humans, thereby increasing the average numbers of genes in the pairwise intersections up to several hundred (Table 1). We found that the conclusions are qualitatively insensitive, with high correlations (0.57 (p value: 0.054) to 0.71 (p value: 0.023)) between host and parasite phases exhibited across the tested ranges of periodicity and activity thresholds (Table 2).

**Table 1.** Gene Intersection Statistics: The average sizes of the pairwise intersections and the average circular variances of pairwise phase shift estimates of periodic genes across all pairs of samples. Interquartile ranges are given in parentheses. Periodic genes are taken to be those whose maximum expression across the time series is at least the specified amplitude threshold in TPMs and which exhibit at least the same degree of periodicity as the top X% of genes when ranked by the JTK-Cycle periodicity scores. Pairs of periodicity thresholds specify those used for humans and *P. vivax* samples.

| Threshold |  | Gene Intersection Size |  | Phase Shift Variance |  |
| --- | --- | --- | --- | --- | --- |
| Amplitude | Periodicity (human, *P. vivax*) | Human | *P. vivax* | Human | *P. vivax* |
| 10 | (3%, 3%) | 7.9 (5) | 45.1 (37) | 2.4e-02 (3.3e-02) | 1.6e-03 (1.3e-03) |
|  | (5%, 5%) | 17.3 (9) | 89.5 (55) | 3.3e-02 (2.4e-02) | 1.9e-03 (1.9e-03) |
|  | (10%, 5%) | 68.8 (28) | 89.5 (55) | 4.0e-02 (2.1e-02) | 1.9e-03 (1.9e-03) |
| 5 | (3%, 3%) | 11.6 (6) | 49.1 (27) | 3.1e-02 (2.6e-02) | 1.6e-03 (1.3e-03) |
|  | (5%,5%) | 30.6 (13) | 93.2 (54) | 3.7e-02 (2.1e-02) | 1.8e-03 (1.8e-03) |
|  | (10%, 5%) | 104.2 (37) | 93.2 (54) | 4.3e-02 (2.7e-02) | 1.8e-03 (1.8e-03) |
| 1 | (3%, 3%) | 22.2 (11) | 49.5 (27) | 3.7e-02 (2.6e-02) | 1.7e-03 (1.6e-03) |
|  | (5%,5%) | 56.9 (27) | 94.1 (53) | 4.1e-02 (2.4e-02) | 1.9e-03 (1.9e-03) |
|  | (10%, 5%) | 207.8 (77) | 94.1 (53) | 4.6e-02 (2.8e-02) | 1.9e-03 (1.9e-03) |

**Table 2.** Phase Circular Correlations: The circular correlations between human circadian phases, *P. vivax* phases, the wall times of the start of each RNA-Seq time series experiment, these wall times adjusted for the self-reported usual participant wake time, and these wall times further adjusted by any self-reported recent perturbation to the usual wake times. Empirical *p* values are given in parentheses. Periodic genes are taken to be those whose maximum expression across the time series is at least the specified amplitude threshold in TPMs and which exhibit at least the same degree of periodicity as the top X% of genes when ranked by the JTK-Cycle periodicity scores. Pairs of periodicity thresholds specify those used for humans and *P. vivax* samples.

| Threshold |  | Phase Circular Correlation |  |  |  |
| --- | --- | --- | --- | --- | --- |
| Amplitude | Periodicity<br>(human, *P. vivax*) | *P. vivax* | Wall Time | Adjusted by<br>Usual Wake | Adjusted by<br>Recent Wake |
| 10 | (3%, 3%) | 0.70 (0.019) | 0.31 (0.178) | 0.23 (0.298) | 0.16 (0.338) |
|  | (5%,5%) | 0.66 (0.037) | 0.20 (0.291) | 0.18 (0.362) | 0.06 (0.430) |
|  | (10%, 5%) | 0.71 (0.022) | 0.02 (0.553) | 0.01 (0.608) | -0.11 (0.594) |
| 5 | (3%, 3%) | 0.59 (0.053) | 0.15 (0.355) | 0.14 (0.407) | 0.04 (0.449) |
|  | (5%,5%) | 0.65 (0.039) | 0.08 (0.450) | 0.03 (0.562) | -0.10 (0.574) |
|  | (10%, 5%) | 0.63 (0.062) | -0.07 (0.648) | -0.10 (0.673) | -0.16 (0.638) |
| 1 | (3%, 3%) | 0.57 (0.054) | 0.18 (0.322) | -0.01 (0.526) | -0.12 (0.612) |
|  | (5%,5%) | 0.60 (0.048) | -0.02 (0.553) | -0.15 (0.666) | -0.23 (0.718) |
|  | (10%, 5%) | 0.62 (0.043) | 0.01 (0.517) | -0.08 (0.623) | -0.14 (0.621) |

## Discussion

This study demonstrates evidence that the transcriptional phase of the *P. vivax* parasite population during the IEC is well aligned with the circadian transcriptional phase of its human host. This study does not address any mechanism(s) that may be driving the synchronization of individual parasites within the population, or the alignment with the host, though several theories have been put forward in other systems (Hawking, 1970; Prior et al., 2018; Reece and Prior, 2018).

This study affirms that *P. vivax* patients can present with 80% trophozoites throughout the day (Figure 1(C) and Table S3) and explains why clinicians should not expect to see fevers spiking at the same time every day in their malaria patients. However, the clinics in which volunteers were enrolled did not operate 24-hour per day, and Study B participants were required to have parasite populations in a specific IEC stage. The combination of these two constraints could, in principle, bias the alignment between host and parasite phases, but only if there were very tight coupling between both the coarse IEC stage and the IEC transcriptional phase, and separately the human transcriptional phase and the time of day. This concern is ameliorated by the fact that no participant was rejected due to their parasites being in the wrong stage and because participant 09’s parasites indicate that the IEC stage inclusion criteria was not highly restrictive to the range of allowable parasite transcriptional phases. Moreover, we observe rather low correlation between the host phase and the time of day, which further diminishes concern about any spurious correlation between the hosts and their parasites caused by study constraints.

The rather weak observed circular correlation between the participant circadian phases and the time of day may have several explanations. Importantly, there exists substantial interindividual variability in the physiological and behavioral manifestations of the circadian system (Roenneberg et al., 2007), (Kalmbach et al., 2017), with the distribution of so-called chronotypes in human populations exhibiting a wide spread—up to 6 hours according to some studies (Refinetti, 2019; Sani et al., 2015). In addition to light as an entrainment signal, behavioral factors that could not be entirely controlled for, such as sleep/wake times, feeding times and exercise are also external cues that can variably influence internal phasing of parts of the circadian system (Buijs et al., 2013). Moreover, the rate and degree of circadian phase adjustment to disruptions or misalignments between behavioral cues and the time of day, as seen in shift workers for example, is highly variable (Gibbs et al., 2002). Interestingly, multiple subjects in the study reported working at night. Finally, we cannot confirm nor rule out the possibility that the parasite populations, through some yet unknown mechanism, are actively perturbing the phase of the host clock to match its IEC phase, as has been seen in other systems (Prior et al., 2019; Rijo-Ferreira et al., 2020).

Across the entire volunteer cohort, we observe significant correlation between the parasite IEC phases and host circadian phases. However, not all participants show the same degree of alignment. One possible explanation, if true, is that coupling between the host and the parasite population is a transient phenomenon that requires some amount of time (following egress from the liver and initialization of the IEC) to become established. From a dynamical systems perspective, this is a very plausible explanation if the alignment of phases has not already been established during the liver stage since the entrainment of coupled oscillators is known to be a transient phenomenon in many biological systems and their models (Abraham et al., 2010; Granada and Herzel, 2009). Interestingly, participant 16, who exhibits one of the largest disparities between host and parasite phases, had no gametocytes present in their parasite population by blood smear microscopy and reported their fever start date on the same day they entered the clinic and the study (Table S3). These observations are consistent with a very recent entry into the IEC and the observation that the concentration of parasites that produce gametocytes increases over the course of infection (Bantuchai et al., 2022). Note that participant 09 also did not show any gametocytemia.

There are multiple limitations of this study. For one, the cohort of participants is small, making it difficult to control for a variety of potentially confounding factors such as age, prior infection, and days since initial infection. Also, transcriptional determinants of IEC and circadian phases from *ex vivo* whole blood measurements are neither known nor expected to be identical for all individuals. Our data-driven solution requires choices of thresholds to define gene lists and, although the conclusions have been shown to hold over a wide range of thresholds, the circadian signals may be corrupted to some extent by the inclusion of irrelevant genes and/or degraded by the omission of relevant genes. We speculate that degradation of the circadian signal could be further exacerbated by the choice to only compute 24-hour periodicity scores of each transcript profile, which is justified by the constraints on the temporal resolution of the RNASeq time series and fact that the free-running circadian periods is known to be highly concentrated around 24 hours across individuals (study participants in (Crowley and Eastman, 2018) exhibited a range between 23.56 to 24.70 hours), unlike the circadian phasing.

The dataset analyzed in this study represents a rich collection of dynamic gene expression data and participant metadata, the depths of which we have only started to explore. Differential expression analysis may enable comparisons between participants and provide hypotheses for correlates with metadata variates. A detailed study of the expression dynamics of participant 09 before and after apparent parasite egress may help distinguish between potential core oscillator components that may be part of an intrinsic oscillator driving rhythmic gene expression and those which may be driven by external cues not present *ex vivo*. Excitingly, gene regulatory network inference tools may be able to elucidate interactions between host molecular circadian cues and parasite IEC to suggest mechanisms of population synchrony and/or parasite-host phase alignment.

Our finding that the parasite IEC appears to be entrained to the host circadian cycle raises two important questions: what is the selective advantage of this alignment and what are the essential entrainment signals? Although there is evidence for increased parasite fitness when parasite IEC and host circadian cycles are aligned in mouse systems (O’Donnell et al., 2011) the mechanism of the advantage is not yet well-established. It has been suggested by studies involving the mouse-infecting *P. chabaudi* parasite that host eating schedules may serve as a mechanism of parasite population synchronization and developmental stage timing (Hirako et al., 2018; O’Donnell et al., 2020; O’Donnell et al., 2022; Prior et al., 2021; Prior et al., 2018). For example, (Hirako et al., 2018) concludes that parasite proliferation occurs during host food intake, and (Prior et al., 2018) suggests that food intake controls the timing of parasite differentiation and schizogony. Finally, by studying malaria synchrony in mouse mutants with disrupted circadian oscillators, (O’Donnell et al., 2020) concludes that malaria parasite replication is most likely coordinated to when nutrients are made available through eating, and not coordinated to the core clock proteins Per1 and Per2.

We cannot confirm nor rule out the role of eating as a population synchronizer or as a phase entrainment signal for *P. vivax* in humans; however, our results partially challenge the generalizability of the conclusions drawn from studies involving mice. First, because *P. vivax* exhibits a 48 hour cycle in humans—unlike *P. chabaudi* which maintains a 24-hour IEC in mice - food intake alone cannot fully account for timings of developmental events during the IEC. Moreover, this study was conducted *ex vivo* over 48 hours, and so the eating schedules of the participants would need to be well represented by the phases of oscillating transcripts in the blood *ex vivo* over the subsequent 48 hours. Since the conclusions in this study are based on transcription dynamics in whole blood, we cannot distinguish between the roles played by peripheral clocks, any blood glucose rhythms, or those of the light-entrained oscillator in the suprachiasmatic nucleus on the phasing of the parasite populations. Finally, although Per1 and Per2 are core clock components in many mammalian tissues, and have been seen to exhibit 24-hour transcriptional rhythmicity in human peripheral blood mononuclear cells (Boivin et al., 2003), they are not seen to be robustly rhythmic in transcriptional profiling of whole blood samples. Moreover, other circadian signals, including non-transcriptional 24-hour oscillations such as peroxiredoxins redox cycles (O’Neill and Reddy, 2011), are also present in human blood and may influence the transcriptional circadian signal we observe in our whole blood samples. Although the peroxiredoxin cycle in humans does not appear to influence the rhythmicity of the malaria IEC (Smith et al., 2020), it is possible that it could influence the phasing of the IEC in whole blood samples.

Discovery of the molecular signals that coordinate the phases of the *P. vivax* IEC and the host circadian rhythm have important implications for new antimalarial therapies. If this coordination between parasite and host rhythms increases the fitness of the parasite in humans as it does in mouse systems, then disruption of these signals could reduce parasite fitness and potentially reduce the severity of disease.

## Acknowledgements

FCM, SBH, and JH were funded by the Defense Advanced Research Projects Agency, D12AP00025. SBH was also funded by NIH grant R01 GM126555-0. KAM gratefully acknowledges funding from the National Science Foundation grant DMS 1847144. We would also like to thank the laboratory and clinical staff at AFRIMS who assisted with execution of the clinical trial (in alphabetical order): Chaiyaporn Chaisatit, Chaiyawat Mathavarat, Chantida Praditpol, David Saunders, Kingkan Pidtana, Kirakarn Kirativanich, Kittijarankon Phontham, Mali Ittiverakul, Mark Fukuda, Norman C. Waters, Paphapas Khonhan, Piyaporn Sai-ngam, Sabaithip Sriwichai, and Watcharintorn Fagnark. We appreciate the support of the Thai Ministry of Health teams at Nachaluai, Buntharik and Ubon Ratchathani. We also appreciate the support of the Office of Disease Prevention and Control 10 and 10.1 (Danai Jaerakul, Chatree Rasribut, Surajit Termwong, Yossunthorn Salabsri, Suwat Dawal, and Supap Silatan), at Vector Borne Disease Control 10.1 Nachaluai (Anichart Santhaweesuk), and at the Buntharik Malaria Clinic (Prasert Surapol and Kreepol Sutawong). Material has been reviewed by the Walter Reed Army Institute of Research. There is no objection to its presentation and/or publication. The opinions or assertions contained herein are the private views of the author, and are not to be construed as official, or as reflecting true views of the Department of the Army or the Department of Defense. The investigators have adhered to the policies for protection of human subjects as prescribed in AR 70–25.

## Author contributions

We adopt the CRediT taxonomy (https://casrai.org/credit/) to categorize author contributions:

Conceptualization: FCM, KAM, IC, JH, SBH
Data curation: FCM, RCM, SAC, IC
Formal analysis: FCM, KAM, RCM, LMS, SAC, IC, SBH Funding acquisition: JH, SBH
Investigation: ARL, MSO, CYC, CMK, LMS, MS, MA, CT1, CT2, KJ, MW, NU, PV, SC, MO
Methodology: FCM, KAM, ARL, CYC, CMK, CT2, MS, KJ, IC, SBH, SC, CC, NU, ND
Project administration: FCM, SBH, IC, MW, PLS
Resources: FCM, PLS, MA, WK, CT1, CT2, KJ, IC, SBH Software: FCM, RCM, CMK, SAC
Supervision: FCM, PLS, MW, NB, MS, WK, IC, SBH, PV Validation: FCM, MA, CT1, WK
Visualization: FCM, RCM
Writing – original draft: FCM, RCM
Writing – review & editing: FCM, RCM, KAM, KJ, IC, MS, SBH, CC, LMS

## Inclusion and diversity statement

We worked to ensure gender balance in the recruitment of human subjects. We worked to ensure ethnic or other types of diversity in the recruitment of human subjects. We worked to ensure that the study questionnaires were prepared in an inclusive way. While citing references scientifically relevant for this work, we also actively worked to promote gender balance in our reference list. The author list of this paper includes contributors from the location where the research was conducted who participated in the data collection, design, analysis, and/or interpretation of the work.

---

**Figure S1.**
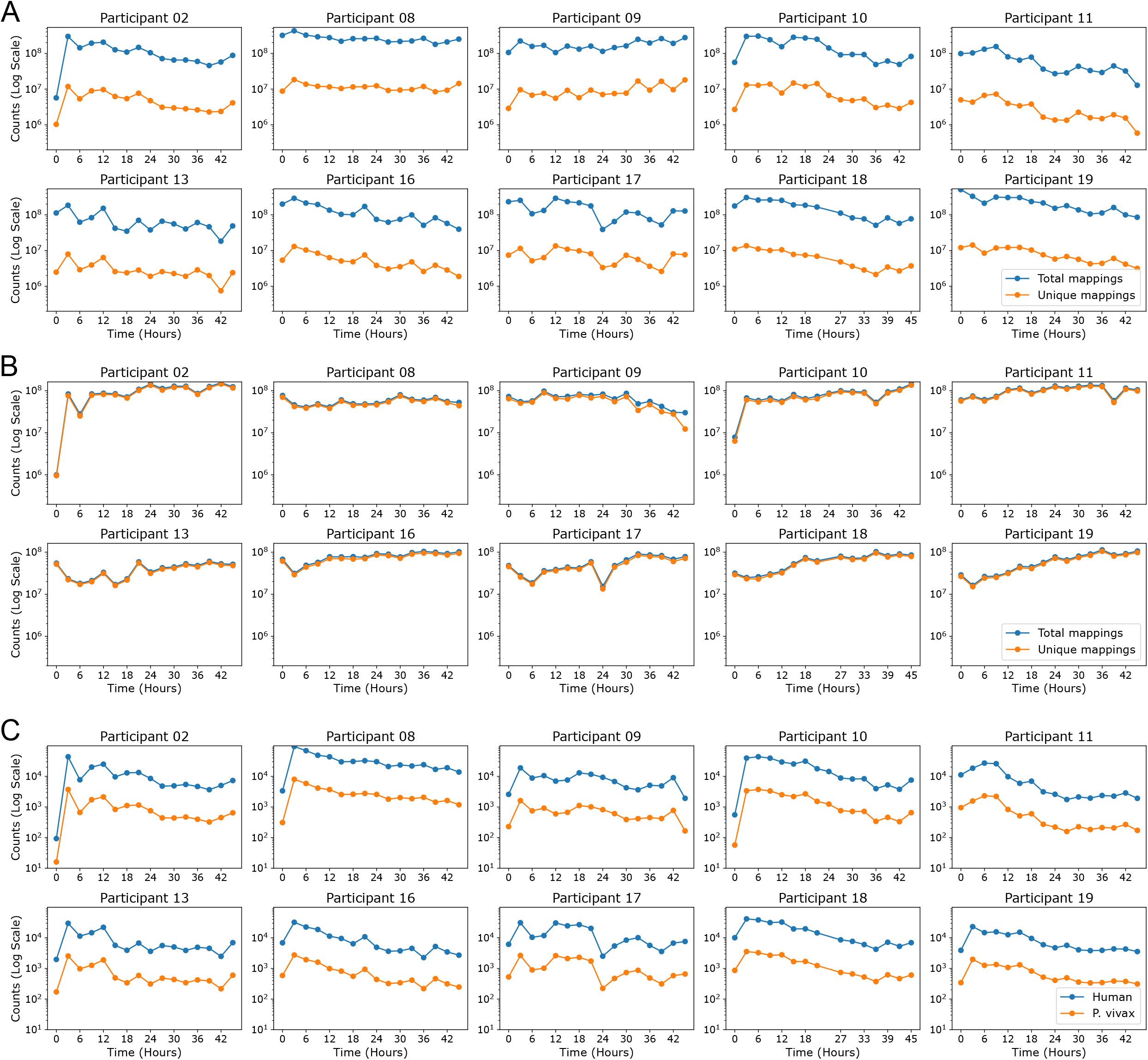
Counts of RNA sequencing read mappings to each organism: (A) For each participant and each time point, the number of mappings of RNA reads to at least one locus, and the number of reads which uniquely map to a single locus in the human transcriptome. (B) For each participant and each time point, the number of mappings of RNA reads to at least one locus, and the number of reads which uniquely map to a single locus in the *P. vivax* transcriptome. (C) For each participant, each time point, and each organism, the number of mappings of RNA reads that were dropped prior to quantification due to reads ambiguously mapping to both human and *P. vivax* transcriptomes.

**Figure S2.**
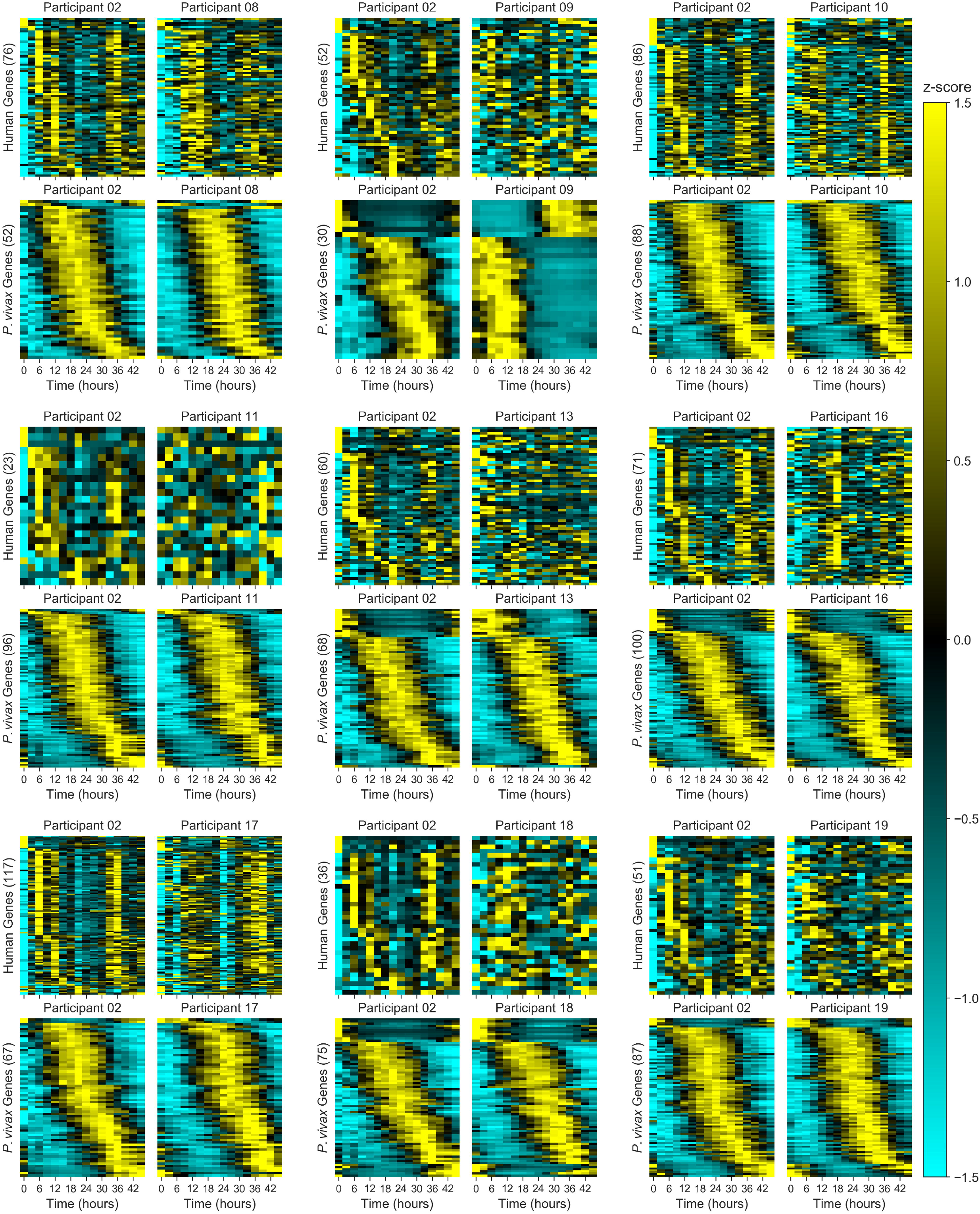
Heatmaps of periodic genes common to participant pairs: Heatmaps of genes which exhibit periodic expression patterns at a specified period---24 hours for human genes and 48 hours for *P. vivax*---in the pairwise intersections of participant 02 and each of the other 9 participants in this study. For each participant and each organism, periodic genes are taken to be those whose maximum expression across the time series is at least 1 TPM and which exhibit at least the same degree of periodicity as the top 5% of genes when ranked by the JTK-Cycle periodicity score. Genes are consistently ordered in each pair by participant 02’s peak expression time during the first period.

**Figure S3.**
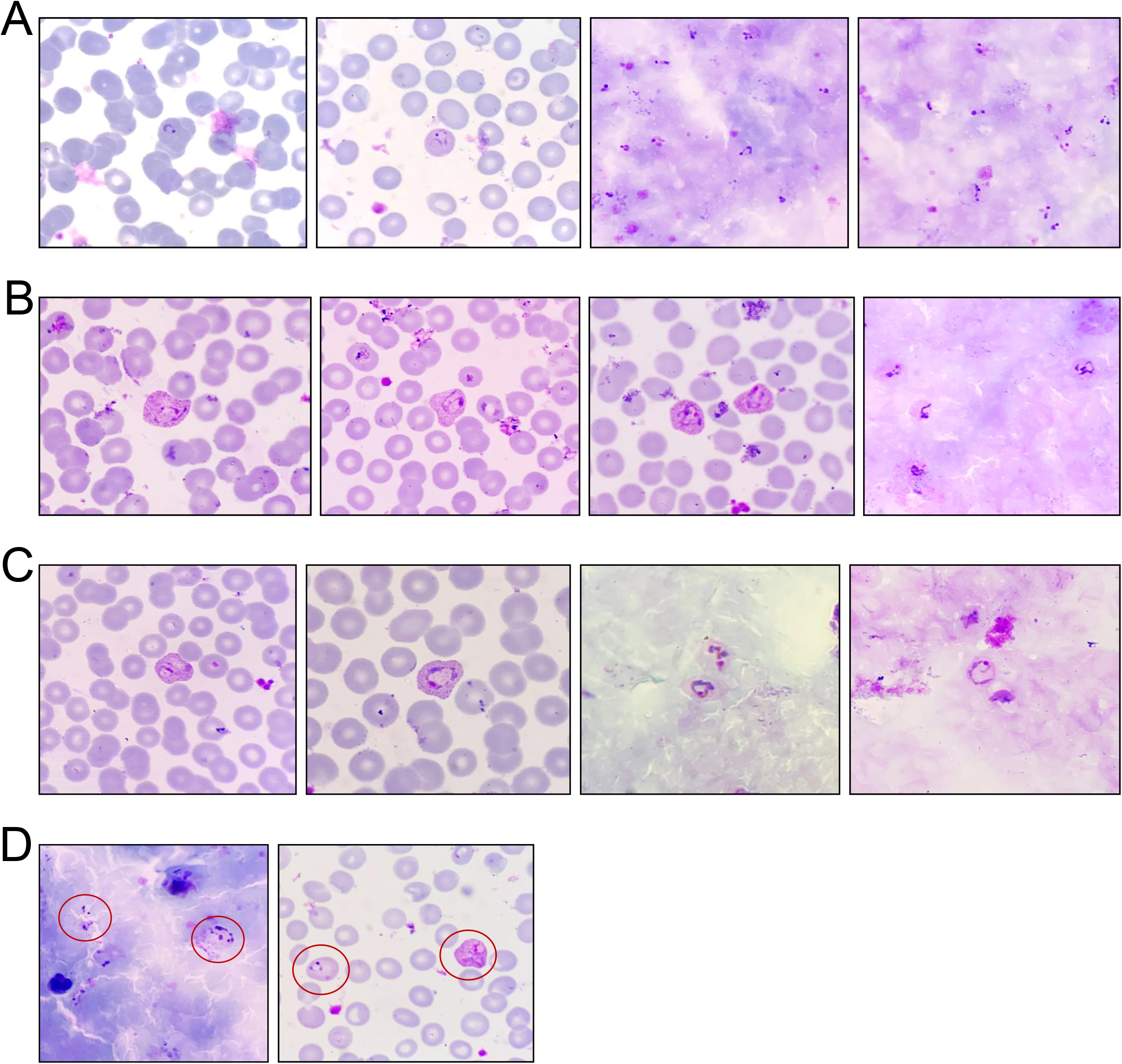
Blood slides of early trophozoite *P. vivax* parasites: (A) Representative examples of *P. vivax* parasites in the beginning of the early trophozoite stage, characterized morphologically by the formation of tiny rings, shown in thin film (columns 1 and 2) and thick film (columns 3 and 4) blood smears. (B) Representative examples of *P. vivax* parasites in the middle of the early trophozoite stage, characterized morphologically by elongated, loose cytoplasm, shown in thin film (columns 1, 2, and 3) and thick film (column 4) blood smears. (C) Representative examples of *P. vivax* parasites in the end of the early trophozoite stage, characterized morphologically by larger and more compact cytoplasm, shown in thin film (columns 1 and 2) and thick film (columns 3 and 4) blood smears. (D) A thick film (column 1) blood smear with parasites in the beginning (left circle) and in the middle (right circle) of early trophozoite stage, and a thin film (column 2) blood smear with parasites in the beginning (left circle) and the end (right circle) of early trophozoite stage.

**Figure S4.**
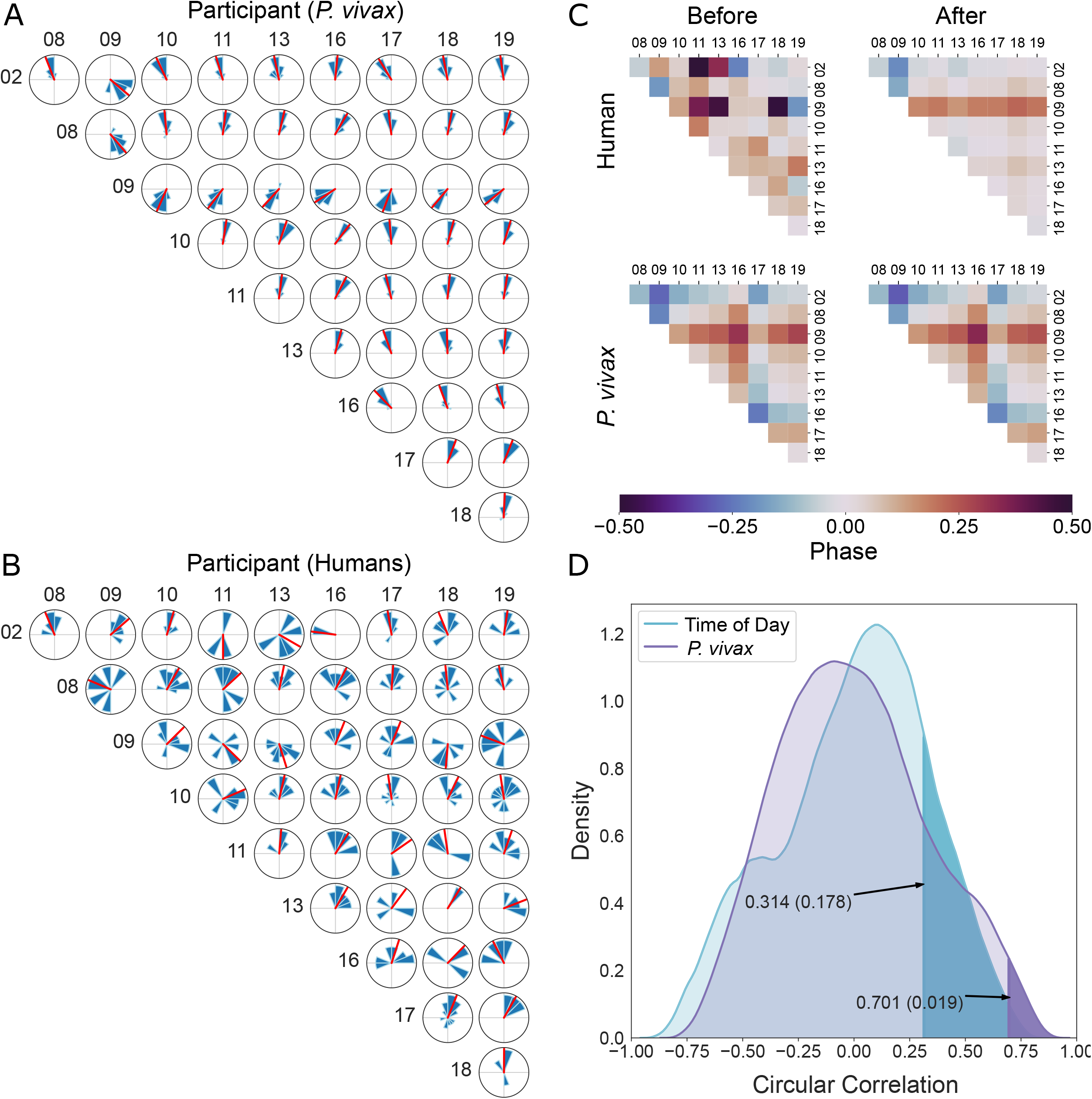
Host/Parasite Phase Correlation Analysis: Circular histograms of phase difference estimates for periodic (A) Human genes and (B) *P. vivax* genes in the pairwise intersection of each pair of participants, and the circular mean phase difference (red radius) of these distributions. For each participant, periodic genes are taken to be those whose maximum expression across the time series is at least 10 TPM and which exhibit at least the same degree of periodicity as the top 3% of genes when ranked by the JTK-Cycle 24/48-hour periodicity score. The radii of the histogram slices have been scaled so the area of each slice is proportional to the number of data points it represents. (C) Heatmaps of the signed phase difference estimates for human participant pairs (top) and *P. vivax* pairs (bottom) before (left) and after (right) phase synchronization was applied within each organism. (D) The empirical distributions of circular correlations between inferred *P. vivax* phases, and time of day (wall times of initial time point in RNASeq experiment), across exhaustive permutations of the inferred human phases. The true circular correlations are indicated with arrows and the empirical *p* values are shown as the darkened regions under the distributions to the right of the true correlations.

## STAR Methods

### Lead contact

Further information and requests for resources and reagents should be directed to and will be fulfilled by the lead contact, Steven B. Haase.

### Materials availability

This study did not generate new unique reagents.

### Data and code availability

- RNASeq data have been deposited at Gene Expression Omnibus and are publicly available as of the date of publication. Accession numbers are listed in the key resources table.
- All original code has been deposited at https://gitlab.com/biochron/2022-human-vivax-coupling and is publicly available as of the date of publication. DOIs are listed in the Key resources table.
- Any additional information required to reanalyze the data reported in this paper is available from the lead contact upon request.

### Experimental Model and Subject Details

#### Study design

This was a minimal risk study enrolling Thai adults diagnosed with *Plasmodium vivax* by rapid diagnostic test (RDT) or microscopy at various medical facilities in Na Chaluai and Buntharik Districts in Ubon Ratchathani Province, Thailand, between September and November, 2016. Those meeting initial inclusion criteria (age >18 years) and exclusion criteria (no signs/symptoms of severe malaria, no antimalarial use in the past 4 weeks and not pregnant (females)), underwent venipuncture for peripheral blood smear and complete blood count (CBC). Smears were read by two trained microscopists to verify parasite species, determine the intra-erythrocyte development cycle (IEC) stage (e.g., percentage early and late trophozoites, and schizonts), and, using CBC results, quantify asexual and sexual parasitemia. Based on blood smear results, the individuals could be enrolled in Part A, Part B or both. Part A, a limit of detection feasibility study, had three enrollment categories: 1. low parasitemia defined as 0.05-0.07% (n=2 subjects), 2. medium parasitemia (0.07-0.19%, n=6 subjects), and 3. high parasitemia (>0.19%, n=2 subjects). Part B subjects, that enrolled for the *ex vivo* time series study, were required to have hematocrit levels that exceeded 25%, a parasitemia level at least 0.1% and the parasite population well-aligned in their IEC stage at early trophozoite only or with the majority (>80%) in the same stage, either early trophozoites or late trophozoites. After the peripheral blood draw, the participants received anti-malarial treatment from Thai Ministry of Public Health staff at the facility to which they presented according to Thai national guidelines. Participant details such as age and sex are available in the attached metadata supplemental information file (Table S3).

All subjects provided written informed consent and the study protocol was approved by the Walter Reed Army Institute of Research Institutional Review Board (IRB #00000794), the Ethical Committee for Human Subject Research Ministry of Public Health, Thailand (IRB #00001629), and the Duke University Institutional Review Board (IRB #00000125).

### Method Details

#### Sample processing & lab procedures

Giemsa-stained thick and thin blood smears were prepared, and asexual and sexual parasitemia independently determined by two microscopists using parasite counts per 200 white blood cells (thick smear) or per 5000 red blood cells (thin smear). Discordant readings were read by a third blinded microscopists and triggered if there was a difference between reader A and B in species, in category assignment for Part A, and for difference greater than 50% for Part A High Parasitemia group (>0.19%) or for Part B. The percentage of the parasites in each IEC stage (early and late trophozoites and schizonts) were determined from total parasite asexual counts obtained during readings.

Part A required 6 ml of blood collected in ethylenediaminetetraacetic acid (EDTA) tubes from which 1ml was used for real-time polymerase chain reaction (PCR) assay. The remaining 5 ml of blood was centrifuged at 2000-3000 revolutions per minute (RPM) for 10±3 minutes and separated into aliquots of plasma and packed cell fraction (buffy coat together with red blood cells), after which all were flash frozen in liquid nitrogen.

Blood drawn for the Part B included 1 ml in EDTA reserved for real-time PCR assay (see the **Real-time PCR detection of malaria** subsection for details), 2 ml of blood in heparin tubes for *ex vivo P. vivax* drug susceptibility testing, and 18 ml of blood in heparin tubes for the *ex vivo* time series experiment. These 18 ml were spun down at 2000-3000 rpm for 10±3 minutes with a plasma layer, approximately 2-4 ml aliquoted, and flash frozen in liquid nitrogen. Of the remaining packed cells fraction (included packed red blood cell and buffy coat layers), 0.2-0.5 ml was placed immediately in liquid nitrogen (time 0-hour sample), and the remaining 3.2 ml was mixed with 156.8 ml of McCoy’s 5A culture media supplemented with 2 mM of L-glutamine, 20% AB serum, generating a 2% haematocrit sample. To collect the culture sample at 3 hour intervals over the subsequent 48 hours, 10 ml of the culture mixture were aliquoted into 25 cm^2^ culture flasks (16 flasks in total for each study participant). Cultures were gassed with 5% CO_2_, 5% O_2_, 90% N_2_ gas and placed in an incubator at 37°C. Approximately every 3 hours, one of the culture flasks was removed, and media and cell pellets were kept in separate tubes and placed immediately in liquid nitrogen, for a total of 17 time point samples including the time 0 sample.

#### Real-time PCR detection of malaria

The whole blood samples (0.2 ml) collected from Part B participants and stored in EDTA tubes were kept frozen at −80°C. Parasite DNA was extracted using the EZ1 DNA blood kit with an automated EZ1 Advanced XL purification system (Qiagen, Hilden, Germany). Multiplex real-time PCR for *Plasmodium* species detection was performed using primers and probes (Kamau et al., 2013; Reller et al., 2013) on Rotor Gene Q 5plex HRM platform (Qiagen, Hilden, Germany). Primers and probes are listed in Table S1. Real-Time PCR reaction was carried out in a total of 25 μl reaction using Rotor-Gene Multiplex PCR kit (Qiagen, Hilden, Germany). Each reaction contained 5 μl of template DNA and a reaction master mix containing 1X rotor gene multiplex master mix buffer, 0.5 μM of each primer and 0.2 μM of each probe. The PCR cycling condition consisted of an initial denaturation step at 95°C for 5 minutes followed by 45 cycles of denaturation at 95°C for 15 seconds and an annealing/extension at 60°C for 15 seconds for *P. falciparum* and *P. vivax* reaction, while the annealing/extension for *P. knowlesi, P. malariae*, and *P. ovale* was conducted at 48°C for 15 seconds. All participants were determined to have *P. vivax* mono-infection.

#### RNA isolation and Sequencing

Frozen cell pellets were used for RNA extraction using the Agencourt RNAdvance Blood RNA purification kit and associated protocols. Samples were cleaned before sequencing with Zymo Research OneStep™ PCR Inhibitor Removal Kit. Isolated RNA was sequenced at the Sequencing and Genomic Technologies Share Resource at Duke University. Libraries were constructed using a kit specific for blood that depleted both ribosomal and globin RNAs. Sequencing was performed on an Illumina HiSeq 4000.

### Quantification and Statistical Analysis

#### Quantitating RNA Sequencing Data

Each participant’s set of Fastq files were aligned to human and parasite genome reference files using STAR (version 2.7.5c) (Dobin et al., 2013) and quantified using RSEM (version 1.3.3) (Li and Dewey, 2011). STAR and RSEM indices were first built using the Gencode v36 reference (reference sequence file GRCh38.primary_assembly.genome.fa and annotation file gencode.v36.primary_assembly.annotation.gtf) for human genome alignment and the PlasmoDB v50 (reference sequence file PlasmoDB-50_PvivaxP01_Genome.fasta and annotation file, PlasmoDB-50_PvivaxP01.gff—which was first converted to a .gtf file using the new code scripts created for this study (Key resources table)—for parasite genome alignment. The STAR command used for building the human STAR index was

~~~
STAR --runThreadN <int> --runMode genomeGenerate --genomeDir <output_directory> --genomeFastaFiles <reference_sequence_file> --sjdbGTFfile <annotation_file>
~~~

The command for building the parasite STAR index was

~~~
STAR --runThreadN <int> --runMode genomeGenerate --genomeSAindexNbases 11 --genomeDir <output_directory> --genomeFastaFiles <reference_sequence_file> -- sjdbGTFfile <annotation_file>
~~~

The RSEM command, which was run once for each organism, used for building the human and parasite RSEM index was

~~~
rsem-prepare-reference --num-threads <int> --gtf <annotation_file> <reference_sequence_file> <output_directory>
~~~

The STAR command for aligning reads from a single Fastq file to the human genome was

~~~
STAR --runThreadN <int> --runMode alignReads --genomeDir <human_STAR_index> --readFilesIn <Fastq_file> --quantMode TranscriptomeSAM --outSAMtype BAM Unsorted --readFilesCommand zcat --outFilterType BySJout --outFileNamePrefix <output_directory> --outFilterIntronMotifs RemoveNoncanonical
~~~

The STAR command for aligning reads from a single Fastq file to the parasite genome was

~~~
STAR runThreadN <int> --runMode alignReads --genomeDir <parasite_STAR_index> --readFilesIn <Fastq_file> --quantMode TranscriptomeSAM --outSAMtype BAM Unsorted --readFilesCommand zcat --outFilterType BySJout --alignIntronMin 10 --alignIntronMax 3000 --outFileNamePrefix <output_directory> --outFilterIntronMotifs RemoveNoncanonical
~~~

Reads from a single Fastq file could map to both organisms’ genomes, which can result in inaccurate quantification results. To remove the small number of reads that map to both organisms’ genomes, we identified reads that were present in both the human- and parasite-associated BAM files generated from aligning a single Fastq file using STAR. These reads were then removed from both BAM files, producing new human- and parasite-associated BAM files that contained reads uniquely mapped to the respective organism. These new BAM files were used in the quantification step. The RSEM command for quantification of the reads uniquely aligned to the human genome was

~~~
rsem-calculate-expression -p <int> --bam --strandedness reverse --fragment-length-mean <avg_fragment_length> --fragment-length-sd <fragment_lenght_std_dev> --no-bam-output <filtered_BAM_file> <human_RSEM_inde×><output_directory>
~~~

The RSEM command for quantification of the reads uniquely aligned to the parasite genome was

~~~
rsem-calculate-expression -p <int> --strandedness reverse --fragment-length-mean <avg_fragment_length> --fragment-length-sd <fragment_lenght_std_dev> -bam --no-bam-output <filtered_BAM_file> <parasite_RSEM_index> <output_directory>
~~~

RSEM produces files containing quantification at the gene-level and the isoform-level, and for each level, produces two types of quantification for each gene from alignments: Fragments Per Kilobase of transcript per Million mapped reads (FPKM) and Transcripts Per Kilobase Million mapped reads (TPM). Gene-level TPM counts were used for subsequent preprocessing.

Alignment and quantification was performed on the Duke Compute Cluster (https://rc.duke.edu/dcc/) at Duke University which, at the time of use, used CentOS 8 as the operating system and SLURM as the scheduler for the entire system.

#### Removal of unexpressed genes

Genes which appear in the comprehensive gene annotation file but which were not expressed (i.e. 0 TPM units) at any time point by a participant or parasite population were removed.

#### Removal of duplicated pseudoautosomal genes

Within the GENCODE comprehensive gene annotation file used for this study are 44 genes with duplicate copies in the pseudoautosomal regions on the X and Y chromosomes (Table S2). Since each of these 44 pairs contained two copies of the same gene, quantification resulted in 44 pairs of duplicated expression profiles for each study participant. Because each participant is male, we added together the duplicate expression profiles in the 44 pairs and retained only a single copy of the duplicated genes. In this way we ensured that pseudoautosomal genes could not have an outsized influence in downstream analyses (e.g. circadian phase estimates) if these genes were found to be highly periodic and sufficiently highly expressed.

#### Quantile normalization

Quantile normalization (QN) was originally developed for normalizing multiple microarray data samples (Bolstad et al., 2003), but is now commonly applied to multiple forms of high-throughput -omics data including RNA-sequencing output (Zhao et al., 2020). In general, the goal is to remove systematic technical variations, including batch effects (Leek et al., 2010) to ensure gene expression levels are more comparable between samples. In this study, species-specific QN was independently applied to each participant’s samples and each participant’s parasite samples. Specifically, genes within each sample (participant, organism, and time point) were sorted according to their TPM values, thereby ranking genes from least expressed to most expressed in each organism and time point. The average TPM value of all genes occupying the same rank in these sorted lists was then computed across all timepoints within each participant and organism and the original TPM value of each gene was replaced with the average of all genes occupying its same rank. Here care was taken to include all equally-ranked genes, including those which are ambiguously ranked because they have identical TPM values within a sample. Finally, each sample was reordered by gene, to realign the now between-sample normalized expression levels.

QN may be regarded as a transformation of gene expression which preserves the relative within-sample expression levels (i.e. the (partial) ranking of genes by their expression is unchanged) while ensuring that the apparent expression of gene X in sample 1 is larger than in sample 2 if and only if the relative abundance (rank) of gene X is larger in sample 1 than its relative abundance in sample 2 (relative to all other genes). This property need not be true for different samples even in TPM units without a judicious choice of between-sample normalization. After QN, the changes in expression level of gene X across time points captures the dynamics of relative abundance of gene X over time.

#### Interpolation of time series expression data

Participant 18’s hour-24 time point sample was accidentally destroyed prior to RNA sequencing. Expression levels for this time point were interpolated following QN using Piecewise Cubic Hermite Interpolating Polynomial (PCHIP) interpolation (Fritsch and Butland, 1984). PCHIP interpolation was also used to up-sample the *P. vivax* microarray expression time series reported in (Bozdech et al., 2008) from 6-hour sampling to 3-hour sampling prior to the time-point correlation analysis presented in Figure 2A. Missing timepoints in (Arnardottir et al., 2014) were also interpolated using PCHIP.

#### Transcript periodicity and phase estimation

Throughout this study, the phase and periodicity of each transcript at a specified period was quantified using the periodicity detection algorithm, JTK-CYCLE (Hughes et al., 2010). For all human and *P. vivax* samples, including those from earlier studies, periods were fixed at 24 or 48 hours respectively. The JTK-CYCLE algorithm compared the time-point ordering determined by the expression levels in a transcript abundance time series to the orderings given by phase-shifted sinusoidal reference curves oscillating at the specified period. Exact *p* values of Kendall’s tau correlation between the experimental abundance profiles and the reference curves were also computed. Each transcript was assigned the phase of the reference curve which minimized Kendall’s tau correlation *p* value (or the average if multiple phase shifts yielded the same *p* value). Each transcript was also assigned the corresponding minimal Bonferroni adjusted *p* value which was used throughout to quantify each transcript’s periodicity at the specified period.

Only transcripts whose QN abundance level exceeded a specified threshold of activity at a minimum of one point during the time series were retained for further analyses, even if a transcript was highly oscillatory according to JTK-CYCLE. Ranges of periodicity and activity thresholds were analyzed to ensure results were insensitive to these choices (Tables 1 and 2).

#### Circadian & IEC phase estimation

To estimate the transcriptional circadian phase, *θ_k_*, of each participant and the transcriptional IEC phase, *ϕ_k_*, of each parasite sample (*k* = 1, …, 10) we first identified transcripts that exceeded specified periodicity and activity thresholds for each of the 10 human samples and separately for each of the 10 *P. vivax* samples. The 45 pairwise intersections of the human transcript lists, and separately the *P. vivax* lists, were computed. In this way we determined, for each pair of participants, the sets of the most oscillatory and most highly expressed human and *P. vivax* transcripts common to that pair. The circular difference between the JTK-CYCLE phase estimates was computed for each transcript in each of the pairwise intersections and the sample circular mean (Jammalamadaka and Sengupta, 2001) was computed for each pair of samples, separately for humans and parasites. More precisely, if *θ*_*m*,1_,…,*θ_m,N_* are the JTK-CYCLE phase estimates (in radians) of participant *m*’s *N* periodic human transcripts–which are also periodic and highly expressed in participant *n*–and *θ*_*n*,1_,…, *θ_n,N_* are the phase estimates for the same transcripts in participant *n*, then the transcript circular phase differences are *δ_k_* = (*θ_i,k_* – *θ_j,k_*) mod 2*π*, for *k* = 1, …, *N*, and the pairwise circadian phase difference estimate between participants *m* and *n* is the circular mean of {*δ*_1_,…, *δ_N_*} given by

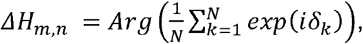

where *Arg*(*z*) is the argument of the complex number *z* = *x* + *iy*, i.e. the angle in [–*π,π*) formed between *z* and the positive real (*x*) axis in the complex plane, and 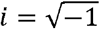. Analogous IEC phase difference estimates, Δ*V_i,j_*, were also computed for each pair of *P. vivax* samples.

Having estimated pairwise circadian phase differences, the eigenvector method for solving the angular (phase) synchronization problem (Singer, 2011) was used to compute globally-consistent transcriptional circadian phases (up to rotation by a constant) for each of the 10 human samples. Likewise, using the pairwise IEC phase difference estimates, we computed globally consistent estimates of the transcriptional IEC phases for each of the 10 *P. vivax* samples. We give a brief summary of this method here. To begin, a 10×10 matrix, **H**, of circadian pairwise phase difference estimates was built, with the *m*-th row *n*-th column being

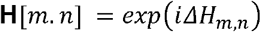

for, 1 ≤ *m,n* ≤ 10. Likewise, the matrix, **V**, encoding the pairwise *P. vivax* IEC phase estimates was built

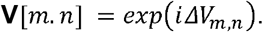

The top eigenvectors, **h** = [***h*_1_**,…, ***h*_10_**] and **v** = [***ν*_1_**, …, ***ν*_10_**], corresponding to the maximal eigenvalues of **H** and **V** respectively, were computed and the circadian phase estimates, 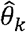, and the IEC phase estimates, 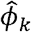, were computed from

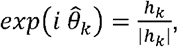

and

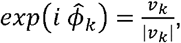

for *k* = 1,… 10. Note, the estimated phases produced by solving the phase synchronization problem are specified only up to a rotation by an unknown fixed angle.

#### Circadian, IEC, & wall phase correlation analysis

To quantify the degree of correlation between the estimated circadian and IEC phases we computed the sample circular correlations (Jammalamadaka and Sarma, 1988) between 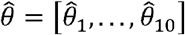 and 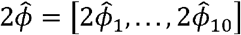 according to the formula

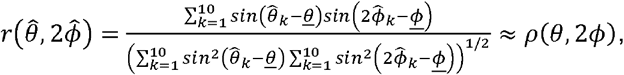

where *<u>θ</u>* is the sample circular mean of 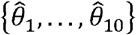 and *<u>ϕ</u>* is the sample circular mean of 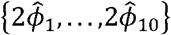 (Table 2), and *ρ*(*θ*, 2*ϕ*) is the exact circular correlation.

Importantly, *ρ*(*θ*, 2*ϕ*) is independent of the choice of the 0-phase for either circular variable, and so is invariant under a rotation of all samples of one of the variables by a constant angle. This is crucial because the solutions to the phase synchronization problem are specified only up to a rotation by an unknown fixed angle. Also, −1 ≤ *ρ*(*θ*, 2*ϕ*) ≤ 1 with *ρ*(*θ*, 2*ϕ*) = 0 if the circular random variables, *θ* and *ϕ*, are independent and *ρ*(*θ*, 2*ϕ*) = 1 if and only if *θ* −2*ϕ* = *c mod* 2*π*, for some constant *c*. Note that if the initiation of the parasite IEC is aligned to a particular host circadian phase, then 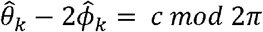, for some constant *c*.

We also computed the circular correlation between the estimated transcriptional circadian phases and the wall-time phases—computed as 2*π t_k_* /24, where *t_k_* is the time of day at the start of the time series RNA-Seq experiment for participant *k*. We further computed circular correlations between the circadian phases and the circular differences between each participant’s time series start time and their self-reported usual wake times. Finally, we computed correlations between the circadian phases and the circular differences between each participant’s self-reported recent wake time, by adjusting the usual wake time by the selfreported recent changes in sleep habits (Table 2 and Table S3).

To measure the significance of the sample circular correlations, we computed empirical p-values through exhaustive permutations of the estimated transcriptional circadian phases. For example, to measure the significance of the correlation between the circadian and IEC phases, we computed the correlations 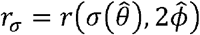 for all 3,628,799 = 10! − 1 (non-trivial) permutations of the sample indices and compared these to the true sample correlation, 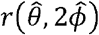. The empirical p-value is then the fraction of permuted sample correlations that are at least as large as the true sample correlation (Table 2 and Figure S4). The same procedure was used to compute the p-values for the other circular correlations reported in Table 2.

## Supplemental information

**Table S1.** Primer and probe sequences for the detection of *Plasmodium spp., P. falciparum, P. vivax, P. knowlesi, P. malariae, P. ovale* and Human RNaseP using multiplex real-time PCR

**Table S2.** Ensemble Id’s of 44 duplicate genes from the pseudoautosomal region contained in the GENCODE comprehensive gene annotation gene transfer format file, gencode.v36.primary_assembly.annotation.gtf

**Table S3**. Table of study Part A and B participant metadata

## Key resources table

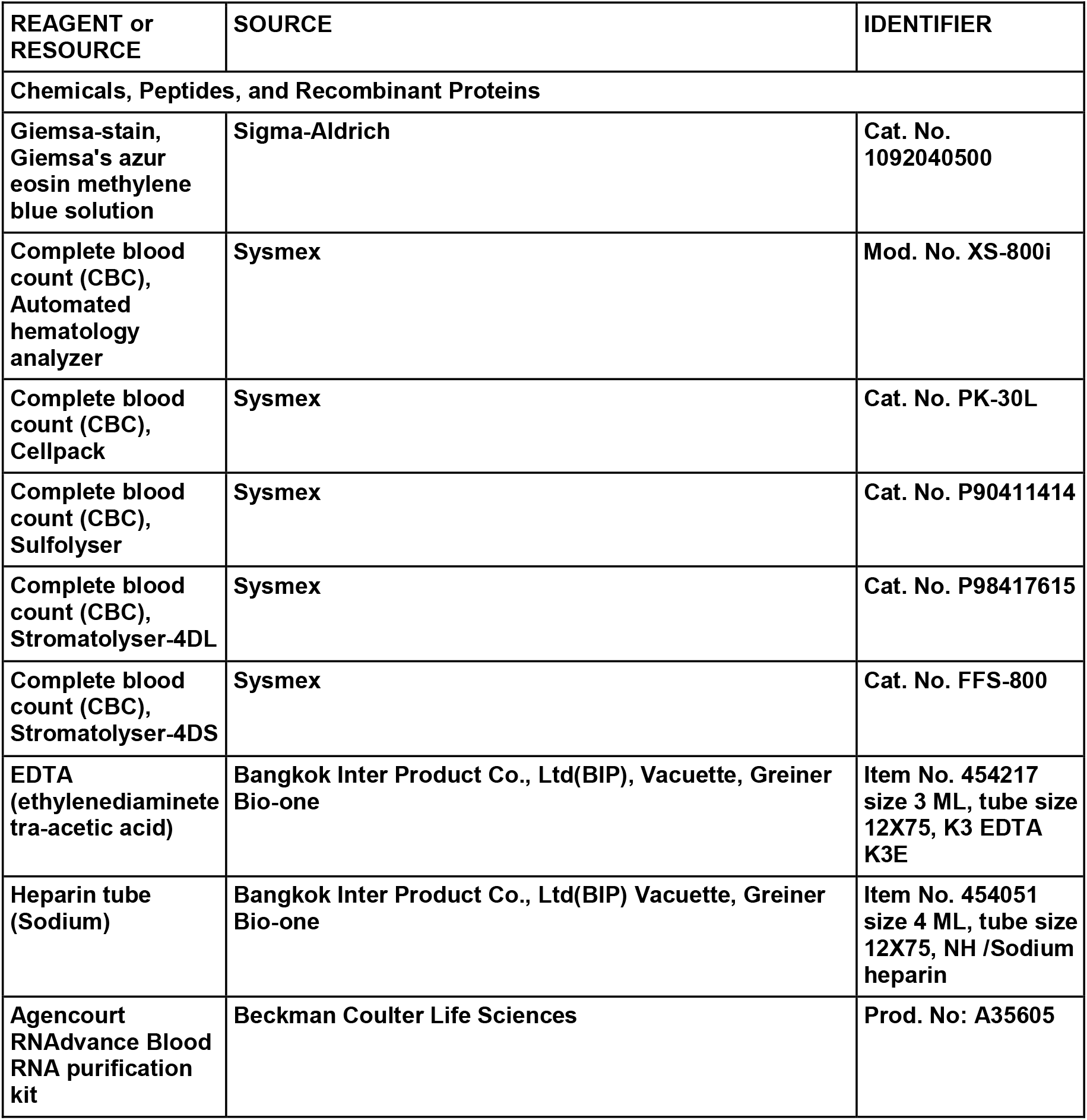
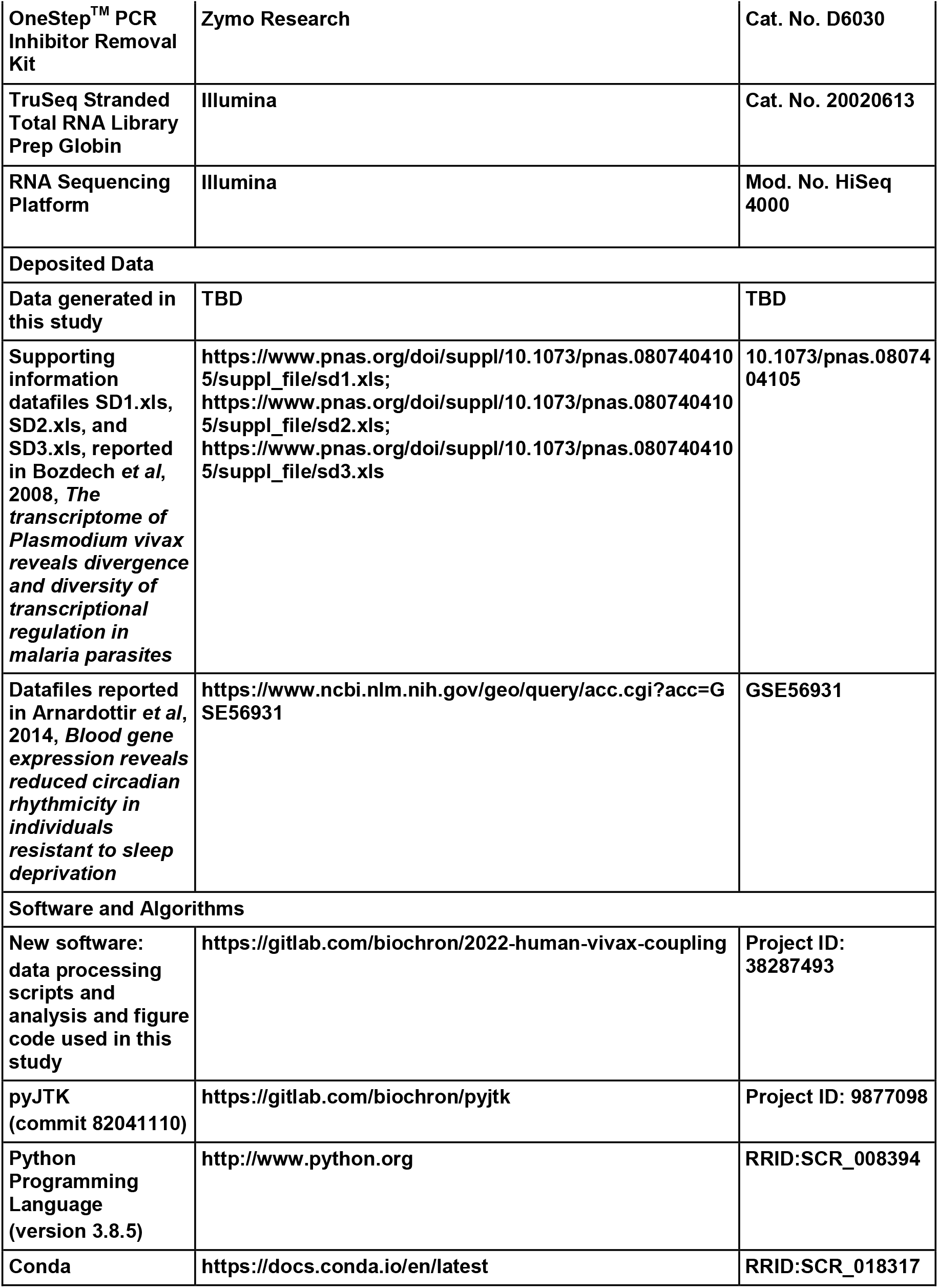
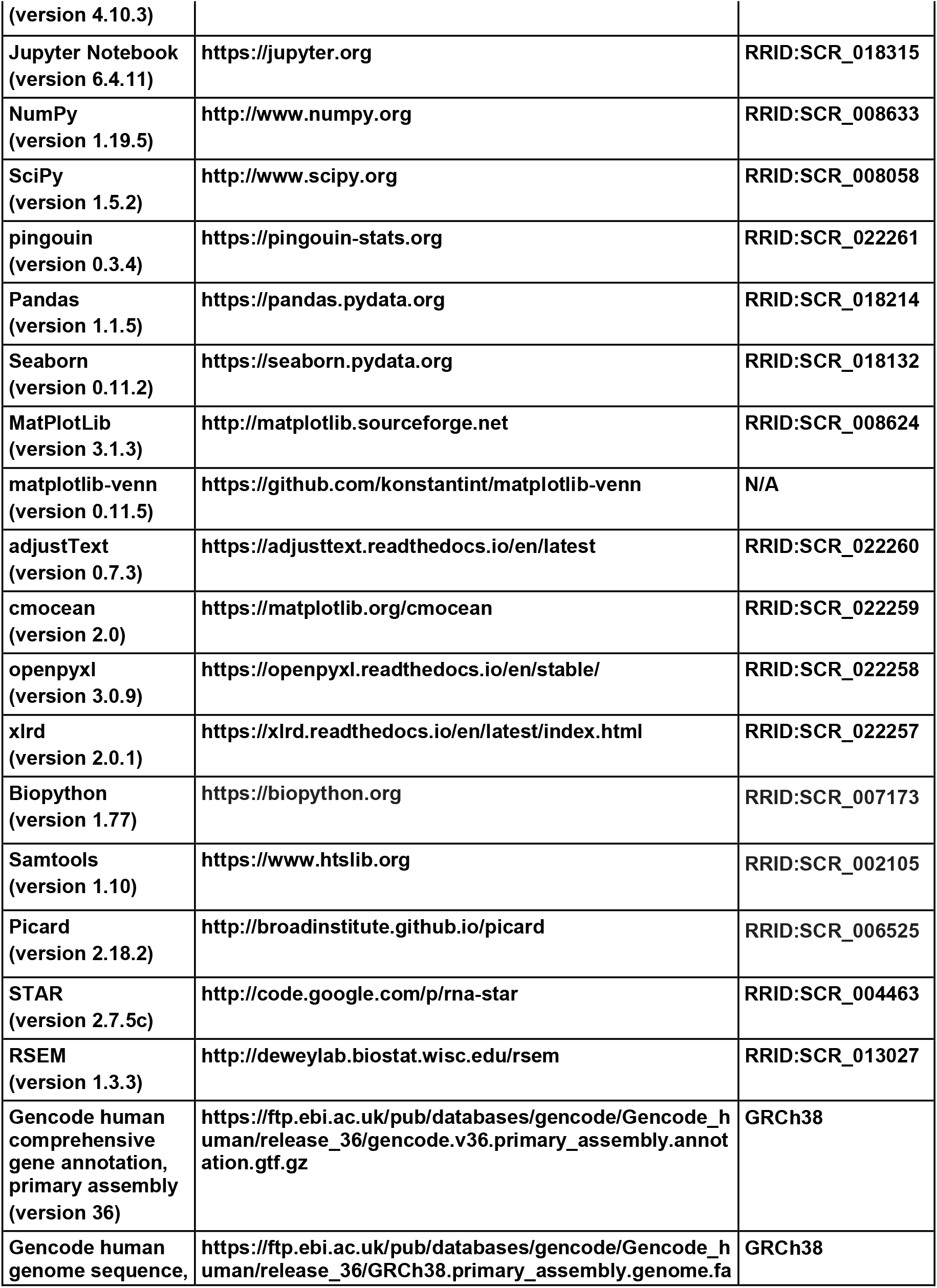
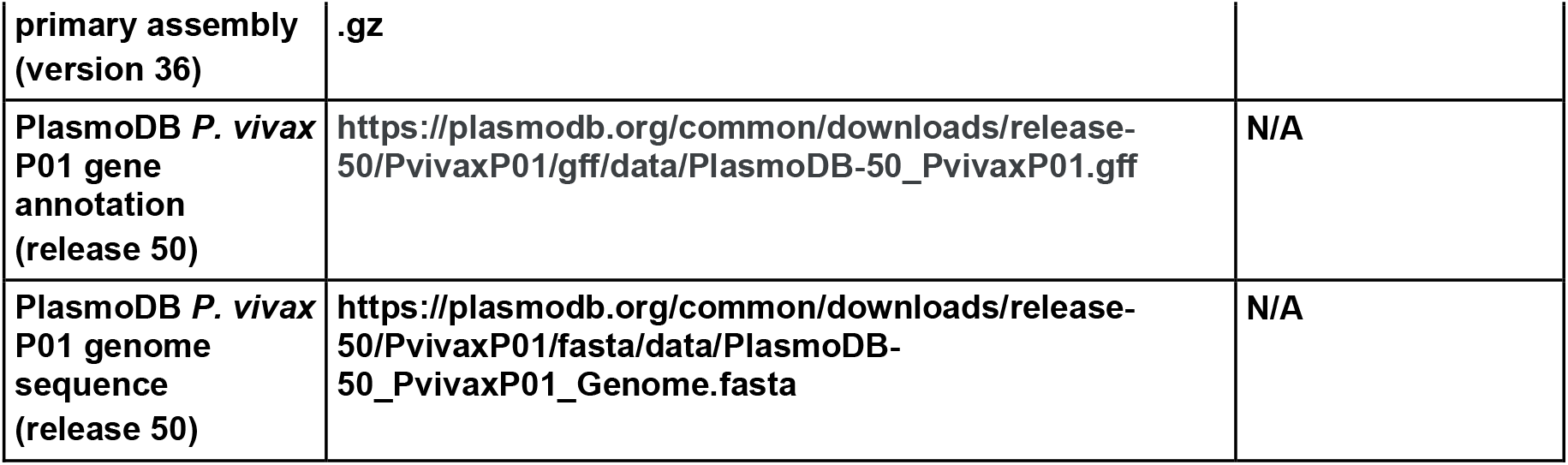

